# Multidimensional encoding of brain connectomes

**DOI:** 10.1101/107607

**Authors:** Cesar F. Caiafa, Franco Pestilli

## Abstract

The ability to map brain networks at the macroscale in living individuals is fundamental in efforts to chart the relation between human behavior, health and disease. We present a framework to encode structural brain connectomes and diffusion-weighted magnetic resonance data into multidimensional arrays (tensors). The framework overcomes current limitations in building connectomes; it prevents information loss by integrating the relation between connectome nodes, edges, fascicles and diffusion data. We demonstrate the utility of the framework for *in vivo* white matter mapping and anatomical computing. The framework reduces dramatically storage requirements for connectome evaluation methods, with up to 40x compression factors. We apply the framework to evaluate 1,980 connectomes, thirteen tractography methods, and three data sets. We describe a general equation to predicts connectome resolution (number of fascicles) given data quality and tractography model parameters. Finally, we provide open-source software implementing the method and data to reproduce the results.

## INTRODUCTION

A fundamental goal of neuroscience is to develop methods to understand how brain networks support function and behavior in individuals across human populations [1–3]. The recent increase in availability of neuroimaging data and large scale projects has the potential to empower new ways of discovery by studying large populations of human brains [4–22]. Exploiting these large-scale data sets will require advances in measurement, computational approaches and theories [23].

Innovation in measurement and computational methods for human brain mapping is shifting the *in vivo* study of the white matter and large-scale brain networks beyond qualitative characterization (such as *camera lucida* drawings), toward structural and functional quantification [24–30]. Tractography and diffusion-weighted magnetic resonance imaging (dMRI) are the primary methods for mapping structural brain connectivity and white matter tissue properties in living human brains. These *in vivo* investigations have shown that there is much to learn about the macrostructural organization of the human brain such that network neuroscience has become one of the fastest-growing fields [3,25,28,29,31–39].

Tractography algorithms use dMRI data to estimate the three-dimensional trajectory of neuronal axons bundles wrapped by myelin sheaths – the white matter fascicles. Fascicles are normally represented as sets of brain coordinates, with coordinates segments spanning anything between 0.01 to 1 mm in length (**Fig. 1a top**). Fascicles have historically been clustered into anatomically cohesive groups called white matter tracts. The largest tracts in the human brain are relatively well characterized and associated with names – such as the corticospinal tract (CST) and the arcuate fasciculus (**Fig. 1b top** [40,41]). White matter tracts communicate between cytoarchitectonically and functionally distinct areas – such as Broca’s or Wernicke’s areas involved in human language processing (**Fig. 1c top** [42–44]). White matter tracts and brain areas together compose a large-scale network called the connectome [45]. Within this network, white-matter tracts represent communication pathways (the edges; **Fig. 1b top**) and brain areas units of information processing (the nodes; **Fig. 1c-top**).

**Figure 1.**
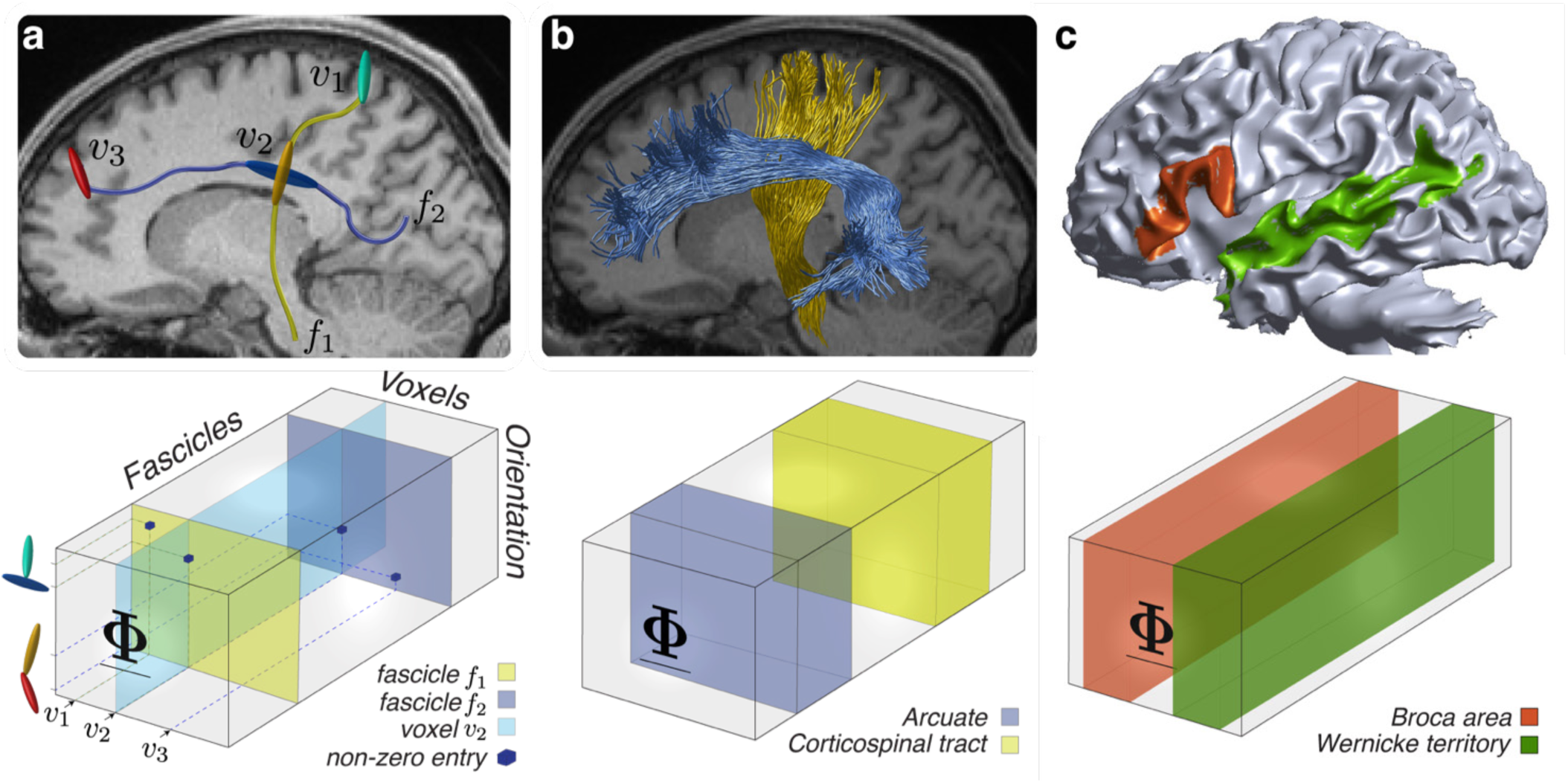
Connectome encoding in tensor space. **(a) Top.** Two white matter fascicles (*f*_1_ and *f*_2_) and three voxels (*υ*_1_,*υ*_2_ and *υ*_3_). **Bottom.** Tensor encoding of fascicles’ spatial and geometrical properties. Non-zero entries in <u>**Φ**</u> indicate fascicles orientation (1^st^ mode), position (voxel, 2^nd^ mode) and identity (3^rd^ mode). **(b) Top.** Two major human white matter tracts (connectome edges). The corticospinal tract and Arcuate fasciculus. **Bottom.** Tensor encoding of connectome edges. The corticospinal tract and Arcuate fasciculus are encoded as collections of frontal slices – blue and yellow subtensors. **(c) Top.** Two human cortical areas (connectome nodes). Wernicke’s territory and Broca’s area. **Bottom.** Tensor encoding of connectome nodes. We show examples of a large temporal area comprising also Wernicke’s territory and Broca’s area encoded as collections of lateral slices – red and green subtensors (areas defined using Freesurfer [42–44,46]).

The standard process to map structural brain connectomes is a threefold *lossy* process. First, dMRI measurements are acquired. Second, tractography is used to identify white matter fascicles. Finally, segmented brain areas are used to identify the terminations of individual fascicles and build a matrix of brain connections. Unfortunately, each of these steps results in loss of information. For example, fascicles are generally estimated using the dMRI data, but after that the data is mostly lost and disregarded in subsequent analyses. Similarly, brain connection matrices are built using the fascicles terminations in segmented brain areas. Yet, once a matrix is built using such terminations, fascicles information is lost; there is no straightforward method to relate back the matrix to the anatomical properties of individual fascicles nor to the dMRI data.

The loss of information during the connectome mapping process, such as described in the examples above, and the lack of frameworks to integrate computations on fascicles, brain areas as well as dMRI data, profoundly limits efforts in clarifying the properties of human brain macroscopic connectivity [2,24,25,47] and white matter microstructure [26,48,49]. Such limitation is especially important because of the established dependency of connectome mapping on brain parcellation schemes and tractography methods [50–57], and the associated need for connectome evaluation methods [50,58,59]. We provide a solution that overcomes information loss in mapping connectomes. The solution has the potential to open new avenues of investigation and fully exploiting opportunities provided by increased data quality and improved tractography methods [60–64]

We propose an integrated connectome encoding framework that prevents information loss during the mapping process. The framework can encode altogether, connectome edges, nodes as well as the associated dMRI data using multidimensional arrays – also called tensors [65–68]. Below we introduce the multidimensional encoding framework and show four applications. First, we use the framework to implement efficiently methods for connectome evaluation. Second, we use the framework to derive a general equation that predicts connectome resolution; namely the number of fascicles supported by data and tractography parameters. To do this we perform a large scale tractography evaluation (13 tracking algorithms, 1,980 brain connectomes, three different data sources [50,61,69]). Finally, we present two additional applications by describing how the framework can be used to perform efficiently statistical inferences on brain connections and white matter tracts using the recently introduced virtual lesion method [50,70] and to chart the reliability and reproducibility in the estimates of the geometrical organization of the human white matter [48,71].

We provide open source software implementing the encoding framework at http://github.com/brain-life/encode, scripts and data to reproduce the analyses in this article at doi:10.5967/K8X63JTX [72,73].

## RESULTS

We present a method to encode the anatomical properties of connectome edges and nodes into multidimensional arrays, also called tensors (see **Methods** [68]). We show an encoding scheme that maps fascicles into the three dimensions of a sparse tensor, <u>**Φ**</u> (**Fig. 1a bottom**). The first dimension of <u>**Φ**</u> encodes fascicles orientation along their trajectory (1^st^ mode). The second dimension encodes spatial position, voxels (2^nd^ mode). The third dimension encodes fascicles indices within the connectome (3^rd^ mode). We show how connectome edges (an ensemble of fascicles) and nodes (an ensemble of voxels) can be conveniently identified in <u>**Φ**</u> subtensors, small subsets of the total volume (see **Fig 1b** and **c**).

Multidimensional encoding of connectomes provides a variety of computational opportunities. This is because direct tensor operations can be applied globally to connectomes. For example, fascicle search, mapping of multiple brain areas and their connections or charting anatomical properties of entire fascicles sets such as their angle of crossing become trivial operands such as finding indices in tensor <u>**Φ**</u>. Below we demonstrate four applications involving such operations.

### First application: Efficient connectome evaluation by tensor encoding

It has been recognized that estimates of brain connectomes can differ substantially depending on the tracking method and data type [48,50,58,71]. Such differences motivated measuring accuracy for brain connectomes in individual brains in order to identify the best fitting connectome model before further studying its properties [50,58].

A few methods to evaluate connectomes and compute errors have been proposed recently [50,76,77]. One of these methods, the Linear Fascicle Evaluation algorithm, or LiFE [50], computes the error of a connectome in predicting the demeaned diffusion signal. LiFE takes as input the set of white-matter fascicles generated using tractography and returns as output the subset of fascicles that predict the dMRI measurements with smallest error (see [50] and **Methods**). LiFE predicts diffusion measurements (vector **y**) in individual brains by combining the diffusion prediction from individual fascicles in a connectome (columns of matrix **M**) as described in equation (7) in **Methods** and **Fig. S2c**. The LiFE model is fit to the data by assigning weights to the fascicles in the connectome (entries in vector **w**; **Fig. S2c**) via a non-negative least-squares method. We show that the LiFE model based on matrix **M**, LiFE_M_ can be accurately approximated using tensor decomposition and the framework introduced in **Fig. 1**.

**Figure 2.**
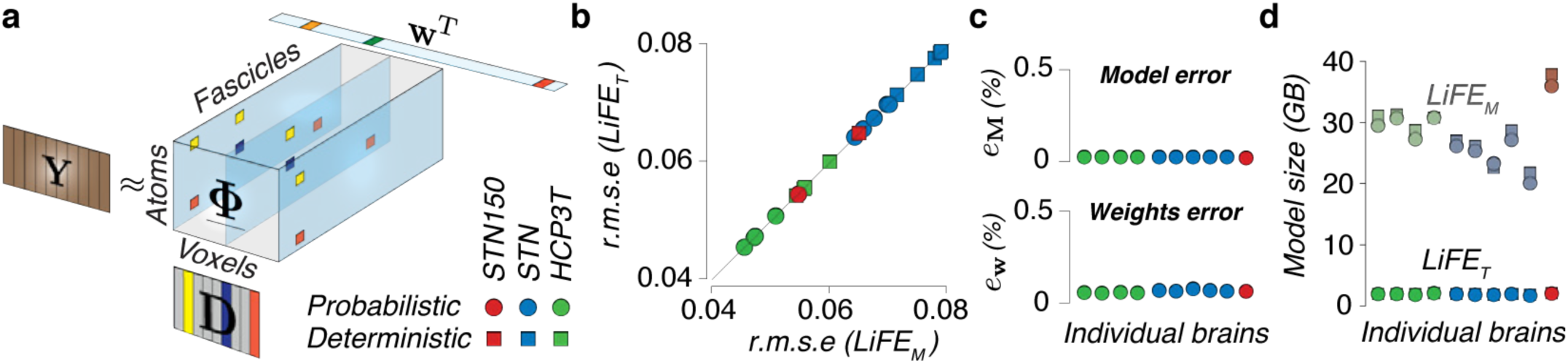
Tensor decomposition of the Linear Fascicle Evaluation method. **(a)**The tensor decomposition model, LiFE_T_(**Y** ≈ <u>**Φ**</u> ×_1_ **D** ×_3_ **w**^T^, see **Supplementary Section 2.1** for details). LiFE_T_ uses a dictionary (**D**) of precomputed diffusion predictions in combination with the sparse tensor, <u>**Φ**</u>, and a vector of fascicles weights (**w**) to model the measured dMRI(matrix **Y**). **(b) Comparison of the error in predicting diffusion.** Scatter plot of the global r.m.s error (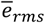; equation (11) **Methods**) in predicting diffusion measurements for LiFE_M_ [50] and LiFET in ten brains, three dataset (HCP3T, STN and STN150 and two tracking methods (tensor-based deterministic and probabilistic tractography). The r.m.s is virtually identical. **(c) Top.** LiFE_T_ error in approximating the LiFE_M_ matrix **Μ**(*e*_M_; equation(12); **Methods**) computed for ten brains(HCP3T, STN and STN150 datasets, probabilistic tractography, *L_max_*=10). **Bottom.** Error (*e_w_*; equation (13); **Methods**) of LiFE_T_ in recovering the fascicle contributions (**w**) assigned by LiFE_M_. (N=10, probabilistic tractography *L_max_*=10) **(d) Model compression.** Measured size of LiFE_M_ (**M**) and the decomposed model, LiFE_T_, (<u>**Φ**</u> and matrix **D**; N=20). Matrices and tensors all stored using double floating-point precision avoiding zero entries [74,75]

**Fig. 2a** depicts the model based on the tensor decomposition, LiFE_T_, where the diffusion measurement (matrix **Y**, equation 23) is factorized into: (1) a dictionary matrix **D** in which each atom (column) represents the precomputed diffusion prediction for a specific fascicle orientation, evaluated at all gradient directions (**θ**, see equation 20), (2) the sparse indicator tensor <u>**Φ**</u> (**Fig. 1c**) and (3) a vector of fascicle weights. **w. Supplementary Results Section 2.1** provides additional details on the decomposition method.

We measured the accuracy of LiFE_T_ in approximating LiFE_M_ using three publicly available data sets: STN, STN150 and HCP3T [50,60,61,78,79]. To do so, we built connectomes in ten individual brains using both, probabilistic (CSD, *L*_*max*_=10 [80,81] and deterministic tractography [82,83], see **Methods**). We report three main results showing that given a sufficient number of dictionary atoms (*L*>360 in; **D**; **Fig. S2d**): (**1**) the global r.m.s. error (equation 11) in predicting diffusion is virtually identical between LiFE_M_ and LIFE_T_ (**Fig. 2b**). (2) LIFE_T_ approximates the LiFE_M_ matrix (**M**) accurately. Specifically, the Frobenius norm-based relative error, *e*_**M**_, is less than 0.1% (**Fig. 2c top**; **Methods**, equation 12). (3) The fascicles weights assigned by LiFE_M_ and LIFE_T_ are virtually identical (**Fig. 2c bottom**, *e*_**w**_ < 0.1%). The relative error between weights estimated by LiFE_M_ and LiFE_T_, *e*_**w**_, was computed using the *ℓ*_2_-norm (**Methods**, equation 13). We show that by increasing decomposition resolution (*L*) the difference in r.m.s., as well as *e*_**M**_ and *e*_**w**_ decrease. See **Fig. S2g**, **h, i, j** and **k** for additional results.

Finally, LiFE_T_ requires a fraction of the memory used by LiFE_M_. To show this, we measured the size of the computer memory used by **M** matrix in the LiFE_M_ model (**Methods**, equation 7) and compared it to that used by **D** and <u>**Φ**</u> together in the LiFE_T_ model (**SI Results**, equation 23). **Fig. 2d** shows measurements in gigabytes for 20 connectomes (500,000 fascicles each, two tracking methods) in ten subjects from the three data sets. Whereas LiFE_M_ can require up to 40GB per connectome, the decomposed model LiFE_T_ requires less than 1GB, a 40x compression factor. All calculations were performed using double precision floating point and sparse data format [74,75]. See Fig. S2 and **Results** for details on the effect of the number of gradient directions (*N*_**θ**_) and connectome fascicles (*N_f_*) on memory consumption.

### Second application: A single equation to predict connectome resolution from data quality

The availability of multiple tracking methods and data types can be both an opportunity or a burden for investigators interested in using them as biomarkers for health and disease [6,7,13,20,21,84]. In an ideal world a single tracking method or data type would supersede all others. Unfortunately, this has been difficult to prove. Not all data and methods are preferable all situations. For example, when measuring patient populations or in developmental or ageing studies might be necessary to measure at lower resolution given time constraints. In principle, higher directional and spatial resolution should be preferred to lower resolution one. Yet, to date we do not have computations to relate data quality and resolution or tractography quality and flexibility to what it is possible to map of the human connectome. Below we defined an equation that predicts connectome resolution (the number of connections supported by the data) given data and tractography quality.

We used LiFE_T_ to perform a large-scale evaluation of the reproducibility of connectome estimates in individual brains to identify the degree to which estimates depend on data quality and tractography. To do so, we generated a total of 1,980 connectomes using thirteen different combinations of tracking methods and parameters on data from twelve individual brains from three sources. Specifically, we used data from (a) HCP3T (4 subjects, 1.25 mm isotropic spatial resolution, 90 diffusion directions [5], (b) HCP7T (4 subjects, 1.05 mm isotropic spatial resolution, 60 diffusion directions [60] and (c) STN (4 subjects, 1.5 mm isotropic spatial resolution, 96 diffusion directions [50].

To test the quality and reproducibility of connectome estimates we generated ten connectomes for each individual brain and tracking method. We used both, probabilistic and deterministic tracking, based on either constrained spherical deconvolution (CSD) or the tensor model [81,82] and generated 500,000 candidate fascicles. We also varied tracking parameters by estimating fiber orientation distribution functions using a range of CSD parameter values (*L*_*max*_= 2, 4, 6, 8, 10, 12). Each one of these 1,980 candidate connectomes was then processed using LiFE_T_. LiFE_T_ identified optimized connectomes, that is, the subset of fascicles with non-zero weight [50] and computed connectomes error in predicting the demeaned diffusion signal (r.m.s.; equation 10). We used this large set of statistically validated, repeated-measures connectomes to test the reproducibility of connectome estimates in individual subjects, as function of tracking method and data type (spatial resolution, signal-to-noise ratio, *SNR*, and number of diffusion directions).

We assessed quality using multiple measures. Connectome quality can be assessed in several ways. For example, the error of the connectome in predicting the diffusion signal can be measured to establish connectome quality [50,76,77]. In addition, connectome resolution, the number of fascicles supported by the data can also inform about connectome quality. Finally, the accuracy of the connectome fascicles can be estimated qualitatively by comparing the anatomical variability of known major white matter tracts estimated from the connectomes using atlases [41]. We established the reproducibility of these three measures across repeated connectome estimates within individual brains and across tracking methods, parameters and data types.

**Fig. 3a** plots mean optimized connectome error and number of found fascicles (±5 standard error of the mean, s.e.m) for the three datasets: STN, HCP3T and HCP7T (1,400 connectomes). The plot shows a series of informative findings. First, data sets naturally cluster into groups, an effect mostly driven by the connectome error, the abscissa. Second, individual brains are separable (along diagonals of the plot) both within and between datasets, such separation is largely independent of tracking method or parameters. Third, for each dataset the number of found fascicles increases with the number of CSD parameters (*L*_*max*_), this is true for both deterministic and probabilistic tracking and can be best appreciated for deterministic tracking, see also inset. Fourth, both the number of found fascicles and connectome error are extremely reliable. LiFE_T_ returns an almost identical number of found fascicles and connectome error across repeated tracking for a given set of parameter and tracking method (error bars are very small compared to the mean values). Finally, results show that probabilistic methods are always better than deterministic methods, showing lower r.m.s. error and higher number of fascicles. This reproduces previous results [50].

**Figure 3.**
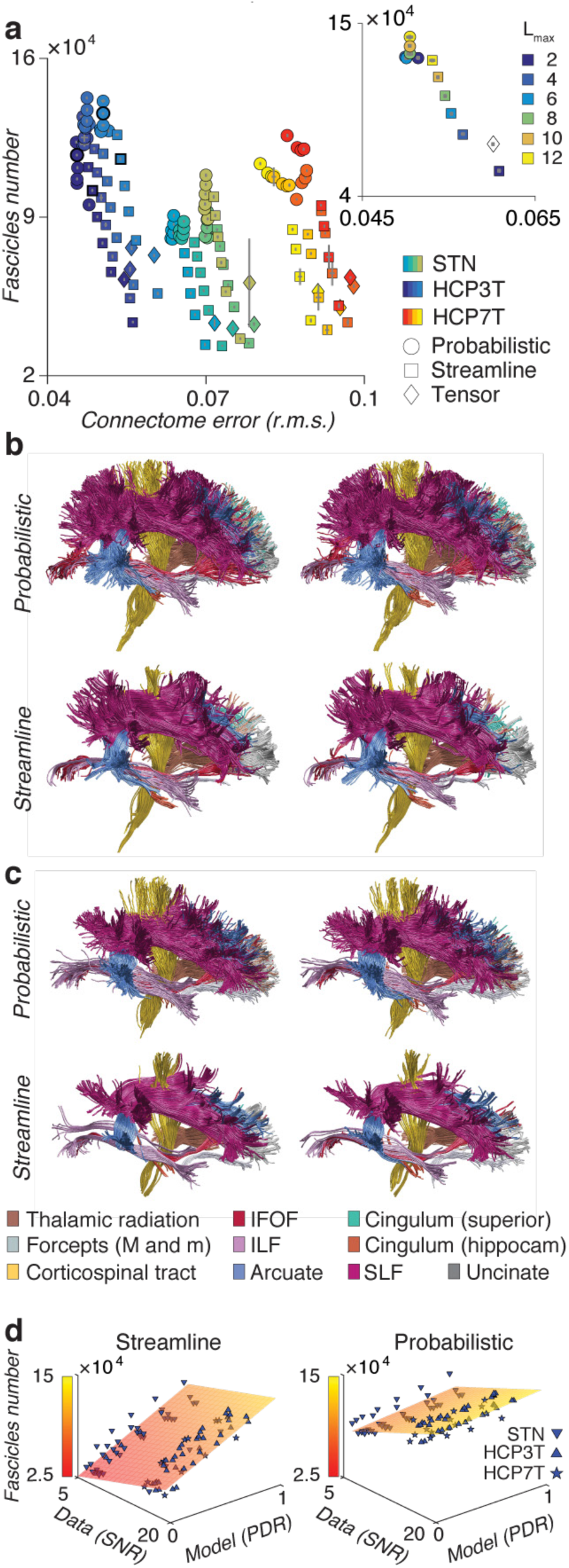
Connectome resolution and anatomical reliability as function of data and method. **(a)** Scatter plot of number of found fascicles and global r.m.s error in LiFE_T_ optimized connectomes (mean ±5) standard error of the mean, s.e.m, N=1,400, n=12 subjects, m=10 repeated tracking, 13, 13 or 9 different *L*_*max*_ values were used for STN, HCP3T and HCP7T, respectively). Inset shows the relation between the number of found fascicles (ordinate) and r.m.s. error (abscissa) and *L*_*max*_ (color) in one subject from the HCP3T dataset. **(b)** Reproducibility of connectome anatomy. Twenty major human white matter tracts, two repeated estimates in a single subject probabilistic (top) and deterministic (bottom) tracking, HCP3T dataset. Tracts anatomy is very similar between repeated estimates when using a single tracking method (compare between columns, top and bottom). Estimated tracts anatomy differs within a single subject when the different tracking methods are used (compare between rows, left or right). **(c)** A different subject from the HCP3T dataset. **(d)** Connectome resolution (fascicles number, *n_f_*) is predicted by data quality (Signal to Noise Ratio, *SNR*) and tractography model quality (*PDR*, number of CSD Parameters to Direction Ratio). Left hand panel, probabilistic tracking(*a*=11,4×10^3^, *b*=2,5×10^3^, *c*=66,3×10^3^, *R*^2^=0.873, p<0.0001). Right hand panel, probabilistic tracking(*a*=6,1×10^3^, *b*=3,4×10^3^, *c*=11,8×10^3^, *R*^2^=0.821, *p*<0.0001) tracking. and are defined in **Methods** equations (2) and (3). **Fig. S3d** shows the marginal distributions of the plots in **(d)**.

We further performed a qualitative evaluation of the degree to which connectomes generated using different tracking methods and optimized with LiFE_T_ show reliable anatomical features. To do so we segmented twenty major human white matter tracts using standard methods and atlases [41,85]. **Fig. 3b** shows two examples of repeated tracts identified in one subject (HCP3T), using probabilistic (top) and deterministic (bottom) tracking. Results show high degree of anatomical similarity for tracts in LiFE_T_ optimized connectomes when using a single tracking method – compare left and right in the top or bottom panels. Conversely, results show anatomical differences within a single individual across tracking parameters –the LiFE_T_ optimization cannot change this result– compare top and bottom tracts. This reproduces previous results [50. **Fig. 3c** shows similar results for a different subject in the HCP3T data set. Importantly, by comparing two different subjects in **Fig. 3b** and **Fig. 3c** it is clearly possible to discriminate between brains based on the anatomical features of the connectomes.

**Fig. S3b** shows multiple examples of major tracts anatomy estimated in individual subjects using repeated connectomes measures. This allows to appreciate the degree of anatomical similarity within subjects given a single tracking method. **Fig. S3c** shows multiple examples of major tracts anatomy estimated in individual subjects using different tracking methods and parameter sets. This allows to appreciate the anatomical differences that different tracking methods generate even within the same subject and data set.

Finally, we identified an equation to predict connectome resolution (number of fascicles *n_f_* in an optimized connectome):

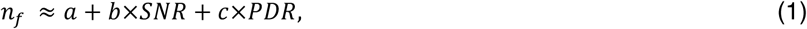

Where data quality is defined as *SNR* = (*m*/*r*)/*V*, *m* and *std* are mean to standard deviation of the non-diffusion measurements, and *V* is the voxel volume (in mm). The tractography method quality was defined as *PDR* = *N_p_*/*N*_θ_, where *N_p_* number of CSD parameters and *N*_θ_ the number of measured diffusion directions (see **Methods,**
equation2 2 and 3)

We used the three data sets (STN, HCP3T and HCP7T) with their different properties –spatial resolution (1.5, 1.25 and 1.05 mm respectively), number of diffusion directions (96, 90 and 60) and *SNR*– and the multiple tractography parameters tested to fit the multilinear model in equation (1). More specifically, **Fig. 3d**, left- and right-hand panels show the connectome resolution estimated in 96 optimized connectomes using different CSD parameters (*L*_*max*_) for all subjects and datasets. Equation (1) was fit to the data using linear regression to estimate, *a*, *b*, and *c*. Equation (1) describes well the fundamental relationship between connectome resolution, data quality (*SNR*) and tractography model parameters (*PDR*). These results provide a first calculation to establish how diffusion data and tractography method affect connectome resolution. In short, connectome density scales linearly with data quality and the flexibility of the tracking method to exploit the data. Equation (1) can inform the choice of dMRI data parameters, such as spatial resolution, directional resolution given data *SNR* and available tracking methods as investigator approach a new study [86,87].

### Third application: Statistical inference on white matter tracts

The concept of virtual lesion has been utilized in several contexts [70,88–91]. More recently, virtual lesions have been used to compute statistical evidence for white matter tracts by measuring the impact of removing entire sets of fascicles from individual whole-brain connectomes [50].

The LiFE_T_ method requires fascicles in an optimized connectome to contribute to the diffusion prediction by assigning non-zero weights, entries of vector **w**, to successful fascicles. Because of this, lesioning fascicles from the model (by setting their weights to zero) increases the prediction error, r.m.s. More specifically, if a set of fascicles, 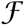, passes through the set of voxels 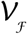, their path-neighborhood, 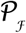, is defined as all fascicles passing through 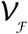 not included in 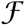. The full signal prediction in 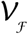 depends on 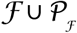. The lesioned model instead, predicts the signal in 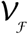 only using 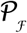. The two models of the signal in 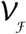, the lesioned 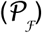 and unlesioned 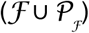 model generate two distributions of r.m.s. error among voxels in 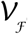. These two distributions can be compared using various measures to establish the statistical evidence for 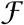 given the data [50].

To date, the virtual lesion method has been employed to establish the statistical evidence for brain tracts and connections [36,38,58,92]. The operations necessary to perform virtual lesions using data represented directly in the brain natural anatomical space require multiple mappings between fascicles coordinates, voxel indices and the corresponding entries in the LiFE model (**M** columns and associated weights). The computational complexity of these operations becomes trivial after encoding connectomes in the tensor framework. We show a visualization of the virtual lesion of the right arcuate fasciculus in a single individual (**Fig. 4a-b**). Given the arcuate fasciculus, 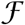 (**Fig. 4a-b**, blue), the identification of 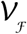 and 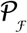 can be achieved in a computationally efficient way using the tensor framework. 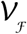 is the set of lateral slices with non-zero entries within the subtensor identified by 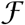 (**Fig. 4b**, yellow) and 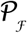 is the set of fascicles (frontal slices) not in 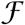 but touching 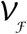 (**Fig. 4B**, red). Computing the signal prediction with and without lesion is then reduced to evaluate the sparse tensor decomposition and consider the tract weights zero (with lesion) or non-zero (without lesion), as shown in **Fig. 4c**.

**Figure 4.**
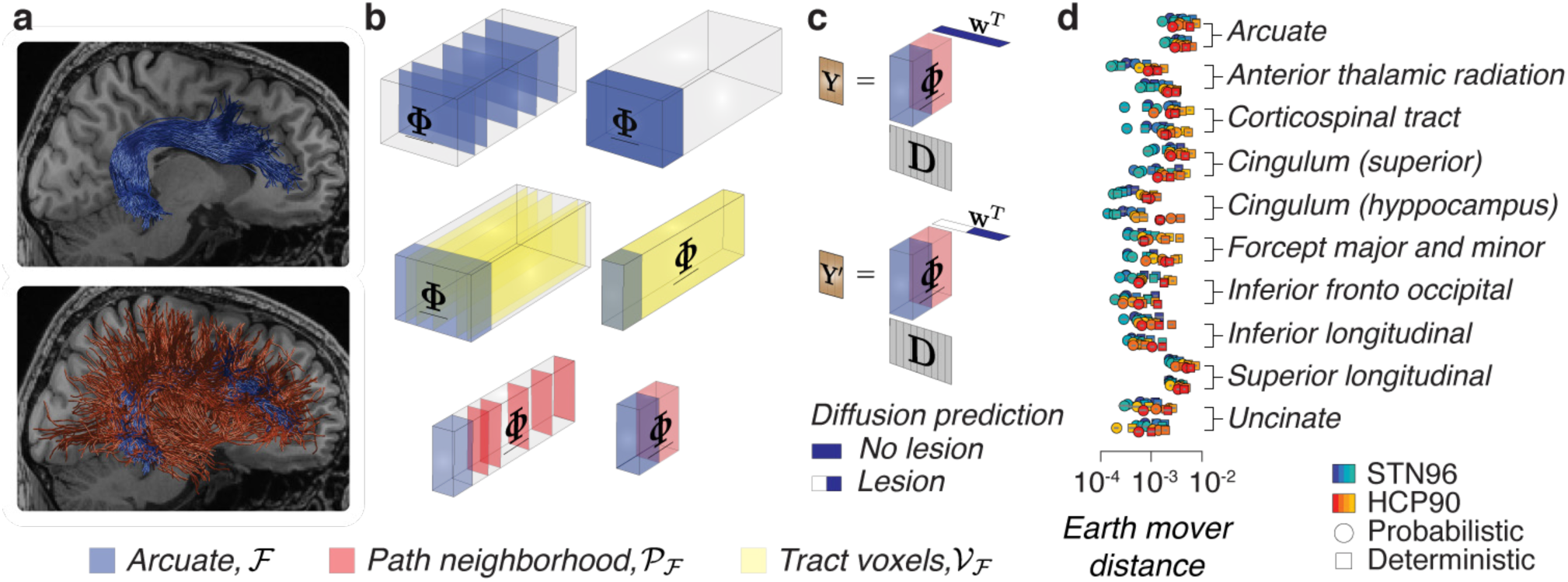
Virtual lesion of white matter tracts using the tensor encoding framework. **(a)** Anatomical representation of the arcuate fasciculus and its path-neighborhood, blue and red respectively. **(b)** Identification of the arcuate fasciculus and its path-neighborhood. Top. arcuate fascicles encoded as frontal slices collated by a permutation 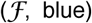. Middle. Ensemble of all voxels touched by the arcuate (lateral tensor slices, yellow) collated by a permutation. Bottom. The path-neighborhood 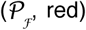 contained in the non-empty frontal slices of 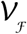. **(c)** The virtual lesion using the tensor framework. Top. Diffusion prediction (**Y**) in the arcuate voxels by the arcuate and its path-neighborhood. Bottom. Diffusion prediction 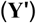 by 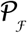, (arcuate fasciculus weights are set to zero, white). **(d)** Statistical evidence for twenty human major white matter tracts [41] established using the sparse tensor encoding framework. Error bars show ±1 s.e.m.

**Fig. 4d** and **Fig. S4** shows the statistical strength of evidence for twenty major human white matter computed with 19,200 virtual lesions (in all connectomes in **Fig. 3**) measured as the earth mover distance [50,93] and strength of evidence [50,93]. These results are important because they reproduce previous findings [50] and show large scale reliability of the *in vivo* statistical evidence of major human white matter tracts validated *post mortem* [40,41].

## Fourth application: Estimates of white matter geometrical organization

Clarifying the geometrical organization of the brain white matter is emerging as an important opportunity given recent improvements in both, measurement and mapping methods [25,26,48,49,94]. Hereafter, we utilize the encoding framework and 160 statistically validated connectomes to quantify the distribution of angles between white matter fascicles associated with pairs of white matter tracts or between tracts and their path-neighborhood [48,71,95].

The corticospinal tract (CST), arcuate fasciculus (Arc) and superior lateral fasciculus (SLF) were segmented in the right and left hemispheres of 160 connectomes estimated using either probabilistic or deterministic tractography in eight brains (STN n=4; HCP3T n=4, *L*_*max*_=10, ten repeated tracking per brain) and standard atlases [41,85]. Angles between pairs of fascicles within a voxel were estimated by operating on the connectome encoding framework (**Fig. 5a-d**). We performed three experiments to establish the dependence of fascicle angles on the tracking method and measured the distribution of angles between fascicles in tracts and neighborhoods. We measured: (a) Crossing angles between fascicles in the Arc and CST at voxels of overlap between the tracts. These fascicles were expected to cross with non-zero degree angle. (b) Angles between fascicles in the Arc and SFL. These fascicles were expected to bypass each other with expected angle near zero degrees. (c) Angles between the Arc and its path-neighborhood. The expected angle of crossing between tracts and path neighborhoods has generated important debates [48,49,71,95].

**Figure 5.**
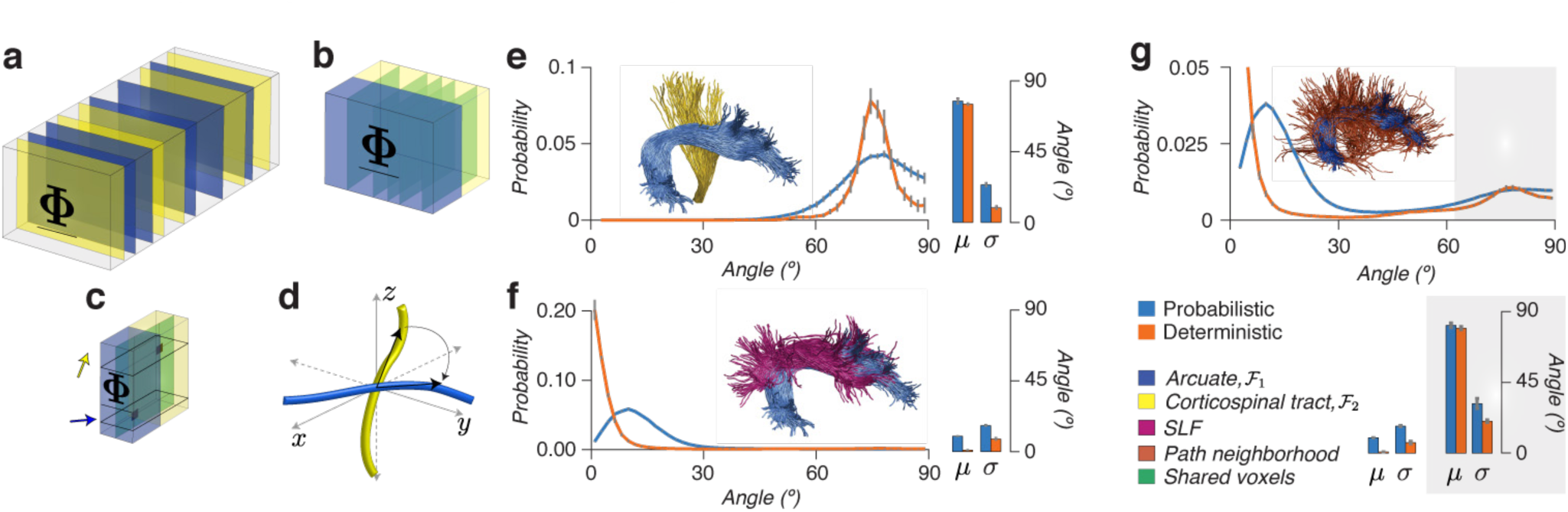
Quantifying variability of estimates for angles of incidence between fascicles in the human white matter. **(a)** Arcuate (Arc, blue) and corticospinal tract (CST, yellow) fascicles identified in frontal slices of <u>**Φ**</u>. **(b)** Voxels shared between Arc and CST located by finding <u>**Φ**</u> lateral slices (green) with non-zero entries in <u>**Φ**</u> subtensors (yellow and blue, respectively). **(c)** Measurement of the angle of incidence in the voxels shared by Arc and CST (green). Angles are determined by finding the indices in the first dimension of <u>**Φ**</u> (1^st^ mode). **(d)** Depiction of angles being computed in brain space. **(e)** Distribution of crossing angles between Arc and CST. **(f)** Distribution of angles incidence between Arc and SLF. **(g)** Distribution of crossing angles between Arc and its neighborhood. Angles computed on Probabilistic (blue) and Deterministic (orange) connectomes (*L*_*max*_=10, STN and HCP3T). Analyses based only on fascicles with positive weight. Histograms show mean across subjects (n=8). Bar plots show peak angle (*μ*) and width-at-half height (*σ*). Error bars ±1 standard error of the mean, s.e.m, across subjects (n=8).

We performed three experiments to measure the dependence of angles between white matter fascicles as function of different tracking methods. In the first experiment, we computed pairwise angles between fascicles associated with either of two tracts, 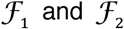, the Arc and CST respectively. We began by identifying the fascicles associated with tracts using the frontal slices of <u>**Φ**</u> (3^rd^ mode; **Fig. 5a**). 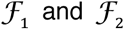 identify two subtensors, **Fig. 5b**, blue and yellow respectively. Voxels containing both 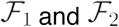 were selected by finding the lateral slices of <u>**Φ**</u> with non-zero entries in both subtensors (**Fig. 5b**, green slices, 2^nd^ mode). Finally, we computed all pairwise angles between fascicles in 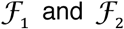 by identifying the atoms (indices in 1^st^ mode) corresponding to the non-zero entries in those lateral slices of <u>**Φ**</u> (**Fig. 5c-d**).

Using the operations described above, we collected distributions of crossing angles, and computed peak distribution (*μ*) as well as width-at-half-max (*σ* **Fig. 5e**). Importantly, we computed approximately 76,000,000 crossing-angles using fascicles validated statistically (fascicles with positive LiFE weight, entries of vector **w**). Crossing angles distributions between Arc and CST peaked approximated at 75º and 78º for deterministic and probabilistic connectomes, respectively (*μ* **Fig. 5e**). The measured *σ* was almost three-fold smaller for deterministic than probabilistic connectomes, 9º and 24º, respectively. These results must be put into context by considering the difference in quality of fit of the two connectomes; where probabilistic connectomes on average have a 4.4% lower error (s.d. 1.4%) and 16.2% higher number of supported fascicle (s.d. 1.1%) than deterministic ones (see Fig. 3a, datasets STN and HCP3T). Fig. S5a shows the same analyses repeated with a different pair of tracts, the CST and SLF. Results are similar for these tracts with distribution peaking (*μ*) approximately at 78.1º and 86.4º for deterministic and probabilistic connectomes, respectively. Measured *σ* was almost two-fold smaller for deterministic than probabilistic connectomes, 17.1º and 31.5º, respectively.

In a second experiment, we measured *μ* and *σ* for the distribution of angles between fascicles within two tracts travelling approximately parallel across the axial plane of the human brain; the Arc and SLF (Fig. 5f). We computed angles distributions for both, probabilistic and deterministic connectomes. The peak distribution (*μ*) was approximately 0º and 15º for deterministic and probabilistic connectomes, respectively. The estimated *σ* were 8.1º and 16.6º, respectively, a 2x increase in variability.

In a final experiment we estimated the distribution of angles between fascicles in a tract, Arc, and its path neighborhood as function of tractography algorithm. Estimates of crossing angles between white matter tracts and path-neighborhoods have been debated [48,71,95]. Hereafter we report *μ* and *σ* for crossing angles between the Arc and its path neighborhood using 8 subjects on STN and HCP3T data sets with probabilistic and deterministic (*L*_*max*_=10) tracking methods. For each subject, we identified the Arc and its path-neighborhood by using tensorial operations similar to the ones described in Fig. 5a-d. Results show characteristic bimodal distributions (Fig. 5g). A majority of the path-neighborhood fascicles show angles between 0º and 20º with tract fascicles (*μ*, 9° and 0° for probabilistic and deterministic tracking, respectively) and around 80° (*μ*, 81° and 80° for probabilistic and deterministic tracking, respectively). The estimated *σ* for *μ* peaking at around 80º were 20.5° and 31.7° for deterministic and probabilistic connectomes, respectively, a 1.5x increase in variability.

Considering that probabilistic connectomes predict the diffusion measurement better than deterministic ones, these results demonstrate substantial variability in the estimates of crossing angles that can be obtained using neuroimaging methods and that the estimates will depend on the data and analysis methods [48,71,95]. This result shows a degree of variability of the estimates consistent with recent reports [30,49].

## DISCUSSION

We presented a connectome encoding framework that provides a solution to the current loss of information problem in modern connectome mapping methods. The encoding framework overcomes information loss by integrating fascicles, edges, nodes and associated dMRI data into multidimensional model. The encoding framework has the potential to empower new ways to study the human connectome by providing investigators an integrated multidimensional relationship between connectome nodes, edges, the anatomy of the white matter fascicles and the associated diffusion-weighted measurements. We provide four applications to show the utility of the connectome encoding framework.

The recent increase in availability, quantity and quality of neuroimaging data and mapping methods poses new opportunities as well as challenges for mapping the human connectome [4–21,96]. dMRI is at the forefront of this data revolution [5,60,96]. Technological advances in dMRI data acquisition have permitted reduction of measurement time by factors up to 8-fold [97–99] and increase in spatial resolution up to 13-fold – when comparing volumetric resolution between clinical and high-field dMRI data [60] (2.5 mm and 1 mm, respectively). Firstly, increased data quality and resolution also means increased size. Secondly, increased availability and diversity of data accompanied by the established variability in results from tractography, makes it difficult to identify a single tracking algorithm, parameter set or data type valid for every study [48,50,58,62,71,87]. For this reason, developing principled methods for evaluating data quality and tractography routinely in their relation to the connectome estimates has become paramount.

The field of connectomics and the study of white matter need improved methods for mapping connectomes in living human brains. To date, best practice in the field to map connectomes is still that of committing to a single tracking method or dataset early on, and then study the results. There are some exceptions to this process [58]. Because the current best practices are limited in what they can achieve, multiple reports have been made highlighting the many methodological limitations as well as the dependency of results on data and algorithms [87,100–104]. As a result, we now understand that no single current tracking method nor data set is likely to solve all reported problems. To date, the fields of connectomics and white matter mapping have been tremendously receptive to issues of validity and reproducibility of results. Criticism is an important aspect fundamentally embedded in the very process of scientific inquiry. We believe that dueling on self-criticism can in the long-run become less effective to advance efforts to map the human connectome. We believe it is of primary importance to focuss our most creative thinking on proposing new, potentially better methods to advance with charting the macroscopic organization of the human connectome.

The concept of routine statistical evaluation of brain connectomes has been recently proposed [50,59,76,77]. The proposal is to build predictive models of the measured dMRI signal from the structure of brain connectomes and compare the model prediction and the data by using statistical methods such as cross-validation [105]. The statistical evaluation approach complements the work on tractography validation based on either synthetic or post-mortem preparations [62,106,107]. Previous work evaluated model accuracy, namely how well a tractography method predicts independent dMRI measurements [50]. The present work advances by measuring model precision, how similar connectome estimates are when using a single tractography method repeatedly.

As number and diversity of datasets increase, statistical evaluation will become a priority for improving brain mapping and results reproducibility [59, 108–115]. We derive a general equation that predicts the number of fascicles in a connectome supported by the data given tractography method parameters and data properties. These results are of interest to investigators planning studies on patient populations or to those developing new magnetic resonance imaging acquisition or preprocessing methods. For example, plots like the ones in **Fig. 3d** can inform on the degree to which for example a new acquisition method would improve on measures of interest of researchers (the number of found connections). Alternatively, investigators interested in measuring children or aging populations might be interested in knowing the connectome resolution achievable given the data they can measure with the available hardware and time constraints.

Tensor decomposition methods help investigators make sense of large multimodal datasets [66,116]. To date these methods have found a few applications in neuroscience, such as performing multi-subjects, clustering and electroencephalography analyses [67,117–122]. Generally, decomposition methods have been used to find compact representations of complex data by estimating the combination of a limited number of common meaningful factors that best fit the data [65,116,123]. We propose a new application. Instead of using tensor decomposition to estimate latent factors and weights, we use a sparse tensor [124] to encode the structure of the brain model; the connectome. This innovative application of tensor decomposition methods in Neuroscience opens new avenues of investigation in mapping brain and behavior using multidimensional and multivariate methods [47,125].

The new application of tensor decomposition proposed here has the potential to allow improving future generations of models of connectomics, tractography evaluation and microstructure [50,58,76,77]. Improving these models will allow going beyond current limitations of the state of the art methods [103]. For example, extensions of the proposed framework would allow building more complex relationships between connectome matrices, edges and nodes without the loss of information of dMRI data and fascicles properties inherent to current methods for connectomics.

We provide an open source implementation of the encoding method and demo files to reproduce figures at http://github.com/brain-life/encode.

## METHODS

### Diffusion-weighted MRI datasets

We use diffusion-weighted Magnetic Resonance Imaging data (dMRI) from three publicly available sources [5,50,60,69].

Dataset are available online at http://purl.stanford.edu/rt034xr8593, http://purl.stanford.edu/ng782rw8378 and https://www.humanconnectome.org/data/.

#### Standord datasets

*STN, 96 gradient directions, 1.5mm isotropic resolution.* dMRI dataset were collected in five male subjects (age 37-39) at the Stanford Center for Cognitive and Neurobiological Imaging using a 3T General Electric Discovery 750 (General Electric Healthcare) equipped with a 32-channel head coil (Nova Medical). dMRI datasets with whole-brain volume coverage were acquired using a dual-spin echo diffusion-weighted sequence. Water-proton diffusion was measured using 96 directions chosen using the electrostatic repulsion algorithm [126]. Diffusion-weighting gradient strength was set to 2,000*s/mm*^*2*^ (TE = 96.8*ms*). Data were acquired at 1.5*mm* isotropic spatial resolution. Individual datasets were acquired twice and averaged in *k-*space (*NEX = 2*). Ten non-diffusion-weighted (*b = 0*) images were acquired at the beginning of each scan. Data acquisition and preprocessing steps are described in [50].

*STN150, 150 gradient directions, 2.0mm isotropic resolution.* dMRI data were acquired in one subject using 150 directions, 2*mm* isotropic spatial resolution and b value of 2,000*s/mm*^2^ (*TE* = 83.1, 93.6, and 106.9*ms*). Data acquisition and preprocessing steps are described in [50].

#### Human Connectome Project datasets

*HCP3T, 90 gradient directions, 1.25mm isotropic resolution.* Data of four subjects, part of the Human Connectome Project, acquired using a Siemens 3T “Connectome” scanner were used. Measurements from the 2,000*s/mm*^*2*^ shell were extracted from the original dataset and used for all analyses. Processing methods are described in [5].

*HCP7T, 60 gradient directions, 1.05mm isotropic resolution*. Four subjects part of the Human Connectome 7-Tesla (7T) dataset were used. Data were collected a Siemens 7T scanner [60]. Measurements from the 2,000*s/mm*^*2*^ shell were extracted from the original data and were used for further analyses.

#### Data quality (Signal-to-noise ratio, SNR)

Data quality was computed as the Signal to Noise Ratio (*SNR*) per units of volume, and is defined as follows:

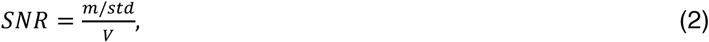

Where *m* and *std* are the mean and the standard deviation of the non-diffusion-weighted signal *S*_0_(*υ*) (see **Supplementary section 1.1**) over all the voxels, and *V* is the volume of a voxel in *mm^3^*. For example, voxel volume for STN dataset is *V_STN_* = (1.5*mm*)^3^ = 3.375*mm*^3^, etc.

#### Whole-brain connectomes generation

Tractography was performed using the MRtrix 0.2 toolbox [81]. White-matter tissue was identified from the cortical segmentation performed on the T1-weighted images and resampled at the resolution of the dMRI data. Only white-matter voxels were used to seed fiber tracking. We used three tracking methods: (i) tensor-based deterministic tracking [81–83], (ii) CSD-based deterministic tracking [80,81], and (iii) CSD-based probabilistic tracking [80,81,127,128]. Maximum harmonic orders (*L*_*max*_) of 2, 4, 6, 8, 10 and 12 were used as long as the number of directions is larger than the number of parameters *N_p_* = 0.5(*L*_*max*_ +1)(*L*_*max*_ +2) [129]. The following parameter values were used for all tracking: step size, 0.2mm; minimum radius of curvature, 1mm; maximum length, 200mm; minimum length, 10mm; and the fibers orientation distribution function (fODF) amplitude cutoff, was set to 0.1.

We created 10 candidate whole-brain connectomes by repeating tracking using 500,000 fascicles in each individual brain dataset (fourteen), tractography method (three) and parameter *L*_*max*_ (six). In addition to the available datasets described above, we simulated new STN and HCP3T datasets with a smaller number of directions (*N_q_* = 60) by eliminating a subset of the gradient directions (choosing the retained directions along surface of a sphere using the electro-static repulsion algorithm [126] as described in [130].

A total number of 1,980 connectomes were generated in this work. For each connectome, fascicles of the twenty major human were identified using Automatic Fiber Quantification – AFQ [85].

#### Tractography model quality (Parameters to Direction Ratio, PDR)

One way to measure the effectiveness of the tractography model to exploit the measured diffusion signal is to compute the ratio of the number of parameters used by the constrained-spherical deconvolution model to identify the fiber-orientation distribution function (fODF). PDR is defined as follows:

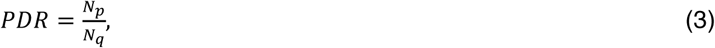

where *N_q_* is the number of gradient directions, for example, the STN dataset has *N_q_*= 96 gradient directions.

#### The Linear Fascicle Evaluation (LiFE) method

Here we introduce the linear model used in [50] to predict diffusion signals based on a multi-compartment voxel model [131,132]. We refer to **Supplementary section 1.1** for an introduction to magnetic resonance diffusion signals.

For a given sensitization strength *b* and gradient direction **θ** the diffusion signal *S*(**θ**, *υ*) measured at a location within a brain (voxel *υ*) can be estimated by using the following equation:

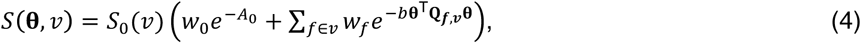

where *f* is the index of the candidate white-matter fascicles within the voxel, *S*(**θ**, *υ*), is the diffusion-weighted signal, *S*_0_(*υ*) is the non diffusion-weighted signal (*b* = 0), *A*_0_ is the isotropic apparent diffusion (diffusion in all directions) and ***Q***_*f,υ*_ is the diffusion tensor matrix (see **Supplementary section 1.1**).

LiFE predicts the demeaned diffusion signal defined as 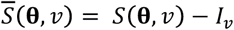, where 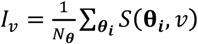 is the mean and *N*_θ_ is the number of gradient directions. Using this definition and equation (4) we arrive at:

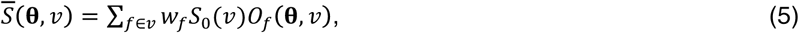

where *0_f_*(***θ***, *υ*) is the orientation distribution function specific to each fascicle, i.e. the anisotropic modulation of the diffusion signal around its mean and it is defined as follows:

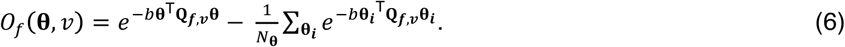

The right-hand side of equation (5) is the prediction model (see Supplementary Fig. 2a-b). The LiFE model extends from the single voxel to all white-matter voxels in the following way (see Supplementary Fig. 2c):

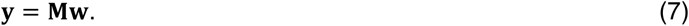

where 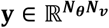 is a vector containing the demeaned signal for all white-matter voxels *υ* and across all gradient directions **θ**, i.e. 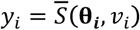. The matrix 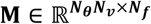 contains at column *f* the signal contribution given by fascicle *f* at all voxels across all gradient directions, i.e.,
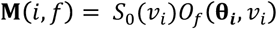, and 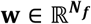 contains the weights for each fascicle in the connectome.

The vector of weights **w** in equation (7) and **Supplementary Fig. 2c** is computed by solving a convex optimization problem [50,76]. More specifically we solve a non-negative least-square (NNLS) problem, defined as follows:

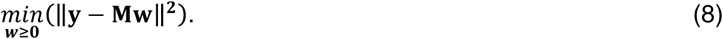

Commonly, the size of the matrix **M** is very large (around 30GB or 40GB for the datasets used here, see **Figure 2d**). Because of this reason, we use NNLS algorithms suitable for large scale problems, such as the BB-NNLS developed in [133].

#### Connectome model prediction error

LiFE predicts the measured (demeaned) diffusion signal using the right-hand side of equation (5). Thus, we can assess the ability of LiFE to model the measured diffusion signal by computing the prediction error in each white-matter voxel. In order to make errors relatively independent of scanner parameters, we compute them on the relative diffusion signal (also referred to as diffusion attenuation), defined as follows:

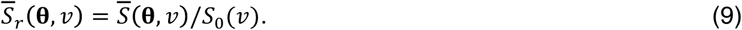

The root mean squared (r.m.s) error in voxel v is defined as follows:

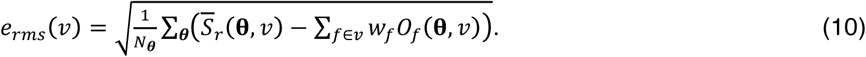

The r.m.s error (equation 10) can be used to compare alternative connectome models. A global r.m.s error 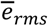 can be computed by averaging *e_rms_*(*υ*) over all voxels:

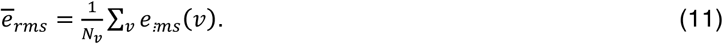

#### LiFE models comparison

We compare a LiFE model matrix **M**(see equation 7) and its approximated version 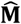 using the relative error:

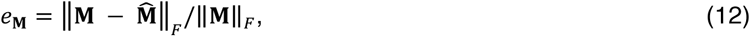

where 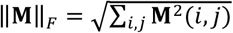 is the Frobenius matrix norm.

Similarly, we compare a vector of LiFE weights and its approximated version 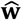 using the relative error defined as follows:

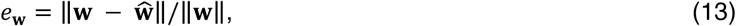

where 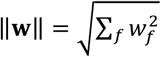 is the Euclidean vector-norm.

#### Tensor notation and definitions

Tensors generalize vectors (1D array) and matrices (2D array) to arrays of higher dimensions, three or more. Such arrays can be used to perform multidimensional factor analysis and decomposition and are of interest to many scientific disciplines [65,66].

Below, we introduce basic concepts and notation (we refer the reader to **Supplementary Table 1**).

*Vectors, matrices and tensors.* Vectors and matrices are denoted using boldface lower- and upper-case letters, respectively. For example, 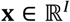 and 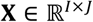 represent a vector and a matrix, respectively. A tensor 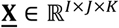 is a 3D array of real numbers whose elements (*i, j, k*) are referred to as **X**(*i, j, k*) or *x_ijk_*. The individual dimensions of a tensor are referred to as modes (1^st^ mode, 2nd mode, and so on).

*Tensor slices and mode-n vectors*. Slices are used to address a tensor along a single dimension and are obtained by fixing the index of one dimension of the tensor while letting the other indices vary. For example, in a 3D tensor 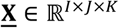, we identify horizontal (*i*), lateral (*j*) and frontal (*k*) slices by holding fixed the corresponding index of each array dimension (see **Supplementary Fig. 1a-top**). Tensors can be also addressed in any dimension by means of mode-*n* vectors. These vectors are obtained by holding all indices fixed except one (**Supplementary Fig. 1a-bottom**).

Subtensors and tensor unfolding. A subset of indices in any mode identifies a volume also referred as to a subtensor. For example, in **Fig. 1d**, we identify a volume by collecting slices in the 3^rd^ mode. In addition, a tensor can be converted into a matrix by re-arranging its entries (unfolding). The mode-*n* unfolded matrix, denoted by
 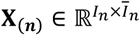, where 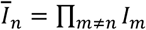 and whose entry at row *i_n_* and column (*i*_1_ − 1)*I*_2_… *I*_*n*−1_ *I*_n+1_… *I_N_* +… (*i*_N−1_ − 1)*I_N_* + *i_N_* is equal to *x*_*i*_1_*i*_2_…*i*_N__. For example, mode-2 unfolding builds the matrix ***X***_(2)_ where its columns are the mode-2 vectors of the tensor and the rows are vectorized versions of the lateral slices, i.e. spanning dimensions with indices *i* and *k* (see **Supplementary Fig. 1b**).

*Tensor by matrix product*. By generalization of matrix multiplication, a tensor can be multiplied by a matrix in a Specific mode, only if their size matches. Given a tensor 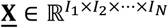 and a matrix 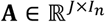, the mode-*n* product

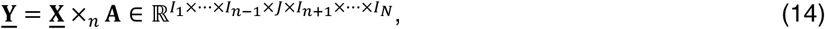

is defined by: 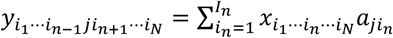, with 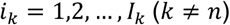 and *j* = 1,2,…,*J*. **Supplementary Fig. 1c** illustrates a 3D tensor by matrix product operation (2^nd^ mode, <u>**Y**</u> = <u>**X**</u> ×_2_ **A**).

*Tucker decomposition.* Low-rank matrix approximation can be generalized to tensors by Tucker decomposition [134]. For example, 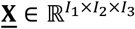, can be approximated by:

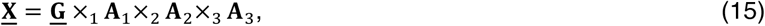

where ×_*n*_ is the mode tensor-by-matrix product. 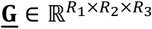 is the core tensor and 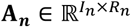 are factor matrices. Such a decomposition guarantees data compression when the core tensor is much smaller than the original, i.e. *R_n_*≪ *I_n_* (see **Supplementary Fig.1d-top**).

*Sparse Decomposition (SD):* Tensors can be approximated also by sparse decomposition [124,135]. In this case, compression can be achieved independently of the size of <u>**G**</u> as long as sparsity is sufficiently high (see **Supplementary Fig.1d-bottom**).

## Author Contributions

F.P. and C.F.C conceived the study. C.F.C. developed the tensor decomposition model. C.F.C. and F.P. designed and performed experiments. C.F.C. and F.P. wrote paper.

## Acknowledgments

This research was supported by (NSF IIS 1636893; NIH ULTTR001108) to F.P. Data provided in part by the Human Connectome Project (NIH 1U54MH091657) and Stanford University (NSF BCS 1228397). We thank O. Sporns, A. Cichocki, E. Garyfallidis, B. McPherson, D. Bullock, S. Ling, M. White, S. Ressl, H. Takemura and B. Wandell for comments and R. Henschel for support.

## 1 Supplementary Methods

### 1.1 Mathematical Modeling of dMRI signals

dMRI measures signals depend on the combination of multiple cellular components within the brain tissue [1] (e.g., neurons, astrocytes and oligodendrocytes). The dMRI signal is generally modeled as the linear combination of two components. One component describes the directional diffusion signal and is presumably related primarily to the direction of the neuronal axons wrapped in myelin sheaths (white matter). This signal is often referred to as *anisotropic diffusion*. The other component describes *isotropic diffusion* (non-directional) and is presumably related to the combination of signals originating from the rest of the cellular bodies within the brain tissue. Below we introduce the equations we used to model the dMRI signal in relation to these components.

dMRI measurements are collected with and without a diffusion sensitization magnetic gradient. Such gradient allows the dMR image intensity to vary depending on water diffusion along a single direction. Generally multiple dMR images are collected for each brain location by varying this diffusion-sensitization gradient (i.e., by sequentially orienting the gradient along several gradient directions). The measured signal depends on a combination of parameters such as diffusion gradient strength and duration. Below we denote the diffusion sensitization gradient strength with the scalar *b* and direction with the unit-norm vector 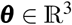.

For a given sensitization strength *b* and diffusion direction ***θ***, the measured dMRI signal at each location within a brain (voxel *v*) can be computed using the following equation [6, 4]:

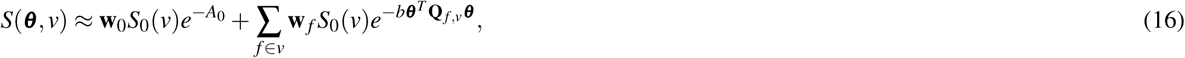

where *f* is the index of the candidate white-matter fascicles in the voxel, *S*_0_(*v*) is the non diffusion-weighted signal in voxel *v* and *A*_0_ is the isotropic apparent diffusion (diffusion in all directions). The value 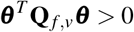 gives us the apparent diffusion at direction ***θ*** generated by fascicle *f*. 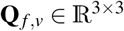 is a symmetric and positive-definite matrix called diffusion tensor [3]. The diffusion tensor **Q**_f,v_ allows a compact representation of the diffusion signal measured with dMRI. Usually, **Q**_f,v_ is represented by an 3D-ellipsoid as shown in **Supplementary Fig. 2a**, which can be mathematically defined with the equation:

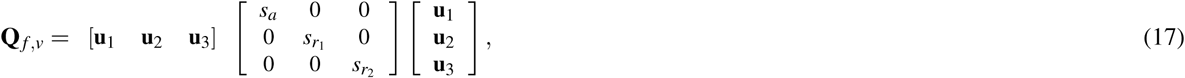

where 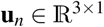 are the unit-norm orthogonal vectors that correspond to the semi-axes of the diffusion tensor ellipsoid, and *s_a_*, *s_r_1__*, *s_r_2__* define the axial and radial diffusivity of the tensor, respectively. In the simplest version of the model, *s_a_* = 1 and *s_r1_* = *s_r_2__* = 0, which means that diffusion is restricted to the main axis direction (stick model) [4,9].

## 2 Supplementary Results

### 2.1 Tensor decomposition of the Linear Fascicle Evaluation model

The LiFE [9] method predicts the demeaned diffusion signal 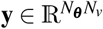 in all voxels (*v* = 1, 2,…,*N_v_*) and gradient directions (***θ***_1_, ***θ***_2_,…, ***θ***_*n*_**θ**__) using the following equation (see **The Linear Fascicle Evaluation (LiFE) method in Online Methods** for its derivation). Each column in matrix **M** (**Methods**, equation 7) contains the diffusion signal contribution from a single fascicle at all voxels and gradient directions. Vector 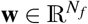 contains the weights associated to each fascicle’s contribution.

Vector **y** and matrix **M** are composed by a vertical concatenation of *N_v_* block vectors 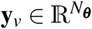 and matrices 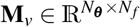, where each block corresponds to a particular voxel *v*(see **Supplementary Fig. 2c**):

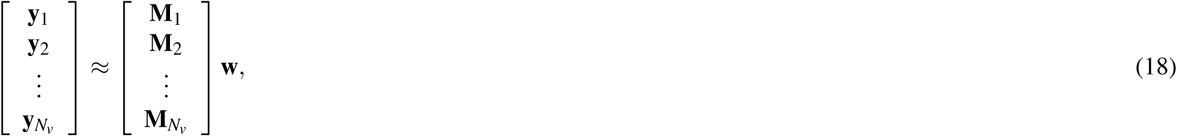

Thus, in each voxel we have the following linear model: *y_v_* ≈ **M_v_w**. Matrix **M**_v_ Can be factorized as follows

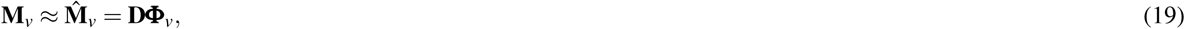

where matrix 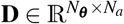 is a dictionary of diffusion predictions whose columns (atoms) correspond to precomputed fascicle orientations, and 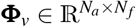 is a sparse matrix whose non-zero entries 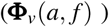 indicate the orientation of fascicle *f* in voxel *v* approximated by atom *a*. The dictionary atoms were created by linearly sampling the azimuth (*α*) and elevation (*β*) of an idealized unit-norm sphere representing the space of putative fascicles directions (*Supplementary Fig. 2f*). More specifically, the entries of the dictionary were computed as follows:

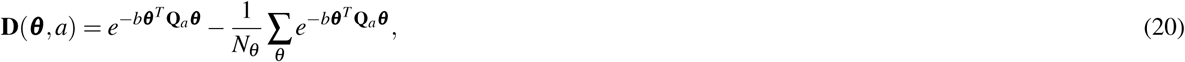

Where **Q**_*a*_ is a diffusion tensor that approximates **Q***_f_,_v_*. At voxels where a fascicle is straight enough, the diffusion signal is approximated by addressing only one atom (one orientation), however for curved fascicles at a voxel its diffusion signal is approximated by a linear combination of few atoms in the dictionary. However, matrix **Φ**_*v*_ follows the constraint

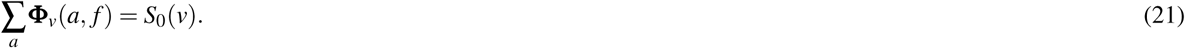

By inserting equation (19) into equation (18), and transforming the approximated full matrix 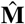 of the original LiFE model [9] into a tensor 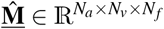 (stacking block matrices **M**_*v*_ as lateral slices^a^), we demonstrate that the following decomposition holds:

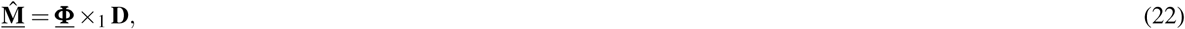

where 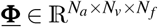 is obtained by stacking all individual voxels matrice 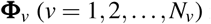 into the lateral slices of <u>**Φ**</u>.

Finally, using equations (7) and (22), the full LiFE_T_ model can be written as (**Fig. 2a**):

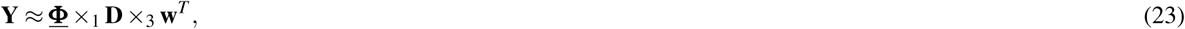

where matrix 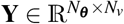 is obtained by stacking vectors *y_v_* (*v* = 1, 2,…,*N_v_*) as columns, 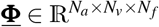 is a sparse core tensor and 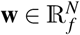 is the vector of weights (see **Supplementary Fig. 2e**). This model is an example of application of Sparse Tucker Decomposition (SD; see **Online Methods**, [5]). In the article we refer to this model as LiFE_T_ (see **Fig. 2a**).

### 2.2 Comparison of the weights estimated by LiFE_M_ and LiFE_T_

In equation (13), we introduced a measure that quantifies the difference between two weights vectors w and 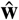. Weights vectors are sparse and their entries indicate which fascicles in the connectome contribute (non-zero) or not (zero) to predict the diffusion signal **Fig. 2c**, bottom panel, shows a global comparison of the vector of weights estimated by LiFE and LiFE_T_ as defined in equation (13).

Hereafter, we perform additional detailed analyses on the difference between these vectors. The observed error between the vectors w and 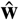 can, in theory, be the result of: (1) each model assigning non-zero weights to a different subset of fascicles; (2) non-zero weights being assigned to the same subset of fascicles but their magnitude differs in the two models; (3) a combination of (1) and (2). This difference is important because it would indicate that either LiFE_M_ and LiFE_T_ select very different fascicles or the same fascicles (2). The case in (1) would indicate a potential bias in LiFE_T_. We explicitly quantified which one of these three cases contributed to the observed error in the weights. To do this, we defined two subsets of weight-fascicles indices. Those that have non-zero values in both models, *common-fascicles indices* (**Supplementary Fig. 2j**, orange) and those that have non-zero values in one model but not in the other, *different-fascicles indices* (**Supplementary Fig. 2j**, green and blue).

We define the vectors of common-fascicles as w*_c_* and 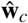 and *different-fascicles* as w*_d_* and 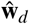 (see **Supplementary Fig. 2j**). We demonstrate that the square of the error of the weights (**Methods**, equation 13) is:

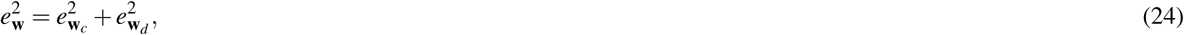

where 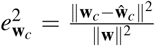 and 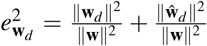 are the squared errors associated to the common and different fascicles, respectively.

### 2.3 Encoding a connectome into LiFE_T_

Encoding a brain connectome information into the LiFE_T_ model involves the computation of the dictionary matrix **D** and the sparse tensor <u>**Φ**</u>.

The dictionary matrix **D** need to be computed first by fixing the total number of atoms *N_a_* to be used, i.e. by setting the minimum incremental unit Δ = *π*/*L* for the spherical coordinates which create a regular grid (see **Supplementary Fig. 2f**). Thus, by increasing the parameter *L* we increase the resolution which has two main consequences: 1) reduces the approximation error (see **section 2.5** and **Supplementary Fig. 2g-i**), and 2) increases the size of the model (see **section 2.6**). We have demonstrated that by using L = 360, we obtained a very accurate approximation and a huge reduction in storage requirements.

By computing the sparse tensor <u>**Φ**</u> we encode the information of fascicles in a connectome into our decomposed model (see **The tensor encoding framework in Results** section). More specifically, the sparse tensor is computed slice by slice (mode-3), i.e. by looking at one fascicle in the connectome at a time. Each fascicle *f* is composed by a set of nodes connected, so for each one of the nodes we need: 1) to identify the voxel index *v* in which the node is located, and 2) find the atom index *a* having the spatial orientation of that fascicle. Finally, for each fascicle node we set a non-zero entry within the sparse tensor <u>**Φ**</u> as follows:

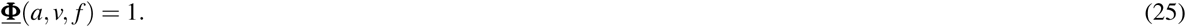

After encoding all the fascicles nodes into the sparse tensor <u>**Φ**</u>, we impose the constraint stated in equation (21) by applying the following normalization:

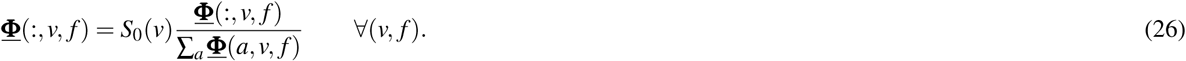

### 2.4 Fitting the LiFE_T_ model

Once the LiFE_T_ has been built, the final step to validate a connectome requires finding the non-negative weights that least-square fit the measured diffusion data. This is a convex optimization problem that can be solved using a variety of Non-Negative Least Squares (NNLS) optimization algorithms. We used a NNLS algorithm based on first-order methods specially designed for large scale problems [8]. Hereafter, we show how to exploit the decomposed LiFE_T_ model in the optimization.

The gradient of the original objective function for the LiFE model can be written as follows:

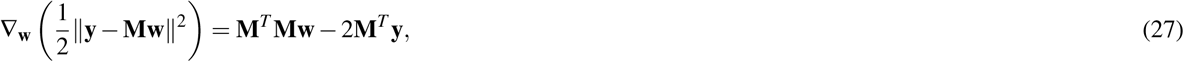

where 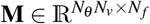 is the original LiFE model, 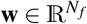 the fascicle weights and 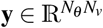 the demeaned diffusion signal. Because the decomposed version does not explicitly store **M**, below we describe how to perform two basic operations (**y** = **Mw** and **w** = **M^T^ y**) using the sparse decomposition.

#### 2.4.1 Computing y = Mw

The product **Mw** can be computed in the following way using a tensor by vector product:

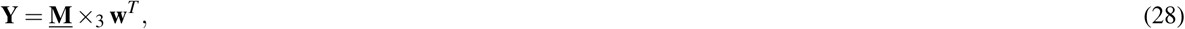

where the result is a matrix 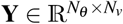, a matrix version of the vector **y**. Using the LiFE_T_ model the product is written as follows:

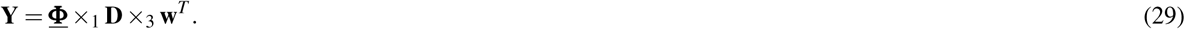

##### Algorithm 1

In Algorithm 1, we present the steps for computing **y** = **Mw** in an efficient way.

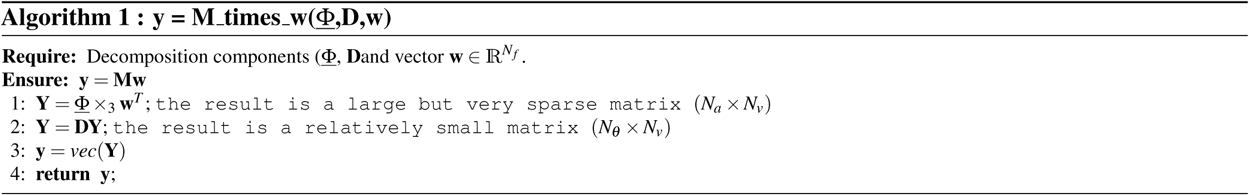

#### 2.4.2 Computing w = M^T^y

The product **w** = **M^T^ y** can be computed using LiFE_T_ in the following way:

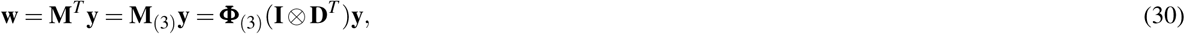

where 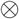 is the Kronecker product and **I** is the (*N_v_* × *N_v_*) identity matrix. Equation (30) can be written also as follows [5]:

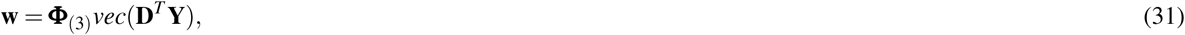
where *vec*() stands for the vectorization operation, i.e. to convert a matrix to a vector by stacking its columns in a long vector.

Because matrix **Φ**_(3)_ is very sparse, we avoid computing the large and dense matrix **D^T^ Y** and compute instead only its blocks that are being multiplied by the non-zero entries in **Φ**_(3)_. This allows maintaining efficient memory usage and limits the necessary number of CPU cycles. In Algorithm 2, we present the steps for computing **w = M^T^ y** in an efficient way.

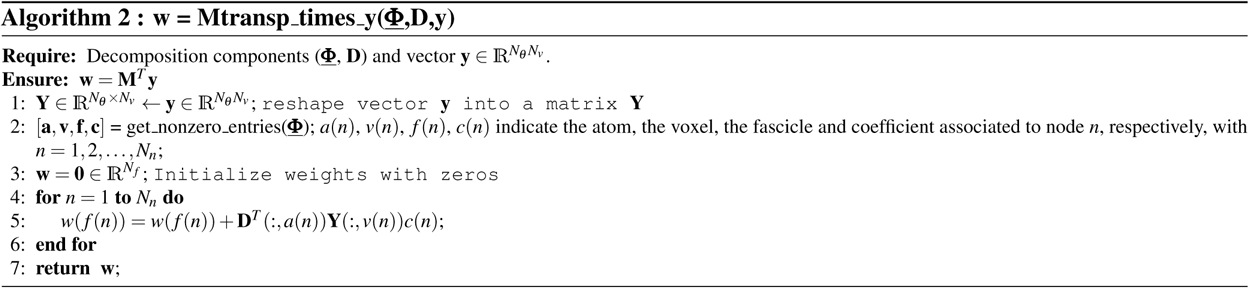

## 2.5 Analysis of the LiFE_T_ model accuracy

The LiFE_T_ model provides an approximation of the original LiFE model. In this section we derive a theoretical upper bound for the model approximation error **e_M_** defined in **Methods**, equation (12) as a function of the discretization parameter *L*, such as 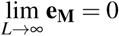

Let us start by focussing in the approximation error of LiFE_T_ model compared to the original LiFE model for a particular voxel *v*, fascicle *f* and gradient direction ***θ***. In this case, the error in modeling the diffusion signal is given by:

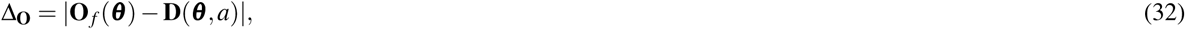

where 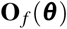 is the diffusion signal as defined in **Methods**, equation (6) (we avoided making reference to the voxel *v* for clearity) and **D**(**θ**; *a*), defined in equation (20), is the diffusion signal of atom a at gradient direction **θ** = [θ_*x*_, θ_*y*_, θ_*z*_]^*T*^. By defining 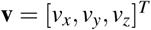 and 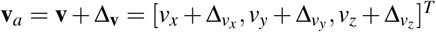 as the vectors pointing out at the directions of the fascicle *f* and its closest dictionary atom *a*, respectively (see **Supplementary Fig. 2f**), and using the “stick” model diffusion tensor, i.e. *s_a_* = 1 and *s*_*r*_1__ = *s*_*r*_2__ = 0 in equation (17), the diffusion tensors of the associated fascicle *f* and its approximation are:

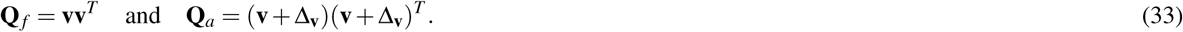

Now, using equations (6), (20) and (33) into equation (32), we arrive at:

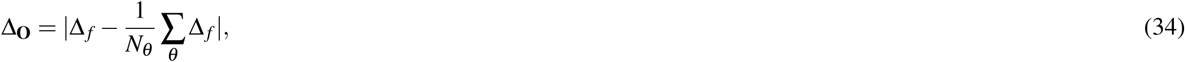

where 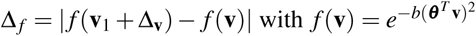.

For a sufficiently small error vector 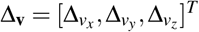, we can approximate Δ_*f*_ as follows:

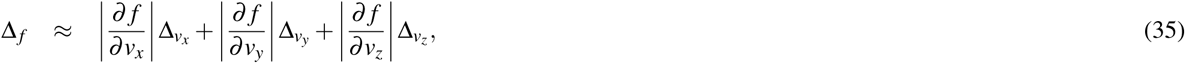

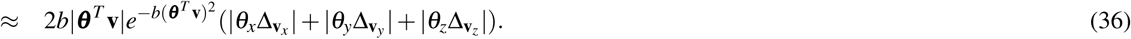

By using the fact that 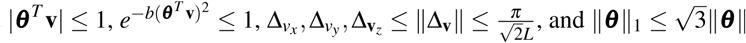, we obtain:

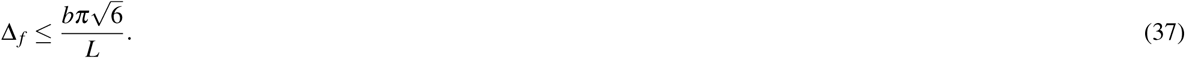

Finally, by using the equation (37) into equation (34), we obtain an upper bound for the error modeling the diffusion signal of one fascicle *f* at one gradient direction in a voxel:

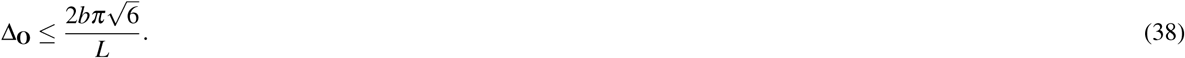

In order to establish a theoretical upper bound of the model error (**e_M_**, **Methods**, equation 12) as a function of the discretization parameter *L*, we need to find an upper bound of
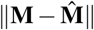 and note ‖**M**‖ is independent of the discretization parameter *L*. The upper bound of 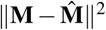 can be obtained by assuming that all fascicles are composed of a fixed number of nodes *N_n_*, and adding up over all nodes *n*, fascicles *f* and directions ***θ*** i.e.

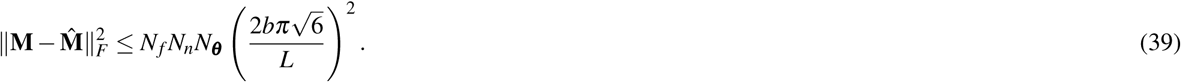

Finally, using equation (39) in equation (12) we obtain the following upper bound of the model error:

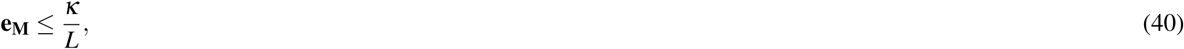

where 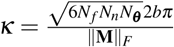 is a constant (independent of discretization parameter *L*).

Equation (40) clearly states that the achievable relative error is inversely proportional to the discretization parameter *L*, which allows us to make the model as accurate as we want by just increasing the dictionary resolution, i.e. increasing *L*.

## 2.6 Analysis of the LiFE_T_ model compression factor

Here, we analytically derive the storage requirements of matrix **M** in LiFE (**Supplementary Fig. 2c**; [9]) and its approximation 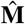 through LiFE_T_, decomposed model (Supplementary Fig. 2d). To do so, as in the previous section, we simplify the analysis by assuming that all fascicles have the same number of nodes *N_n_* and that there are no more than one node per fascicle, per voxel. Under these ideal assumptions the amount of memory necessary to store each fascicle *f* in a sparse matrix **M** is 3*N_**θ**_ N_n_*, since using a sparse matrix structure, three numbers are required for each node, i.e. the row-column indices plus the entry value. Thus the storage cost of *M* is:

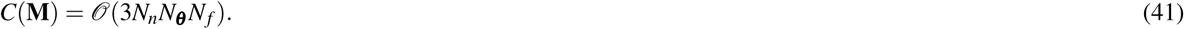

Conversely, storing fascicles in the LiFE_T_ model requires 4*N_n_* values plus the dictionary matrix (i.e. the set of the non-zero entries and their locations within the tensor <u>**Φ**</u> together with matrix **D**). Thus the amount of memory required in LiFE_T_ model is:

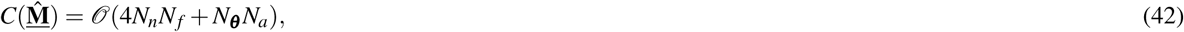

where *N_**θ**_ N_a_* is the storage associated with the dictionary matrix 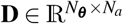. The Compression Factor can be straightforwardly computed as follows:

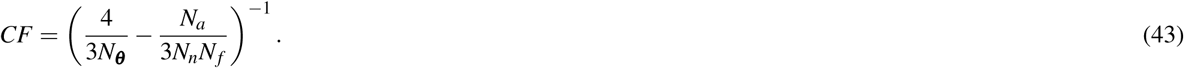

Given that, usually 3*N_**θ**_ N_f_* ≫ *N_a_*, the compression factor can be estimated^b^ as follows:

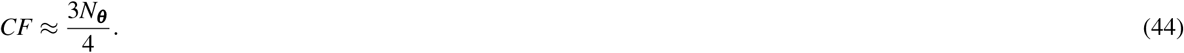

Equation (44) states that the compression factor is proportional to the number of directions *N_**θ**_*, which represents a substantial reduction in memory requirements for modern datasets.

**Supplementary Fig. 1.**
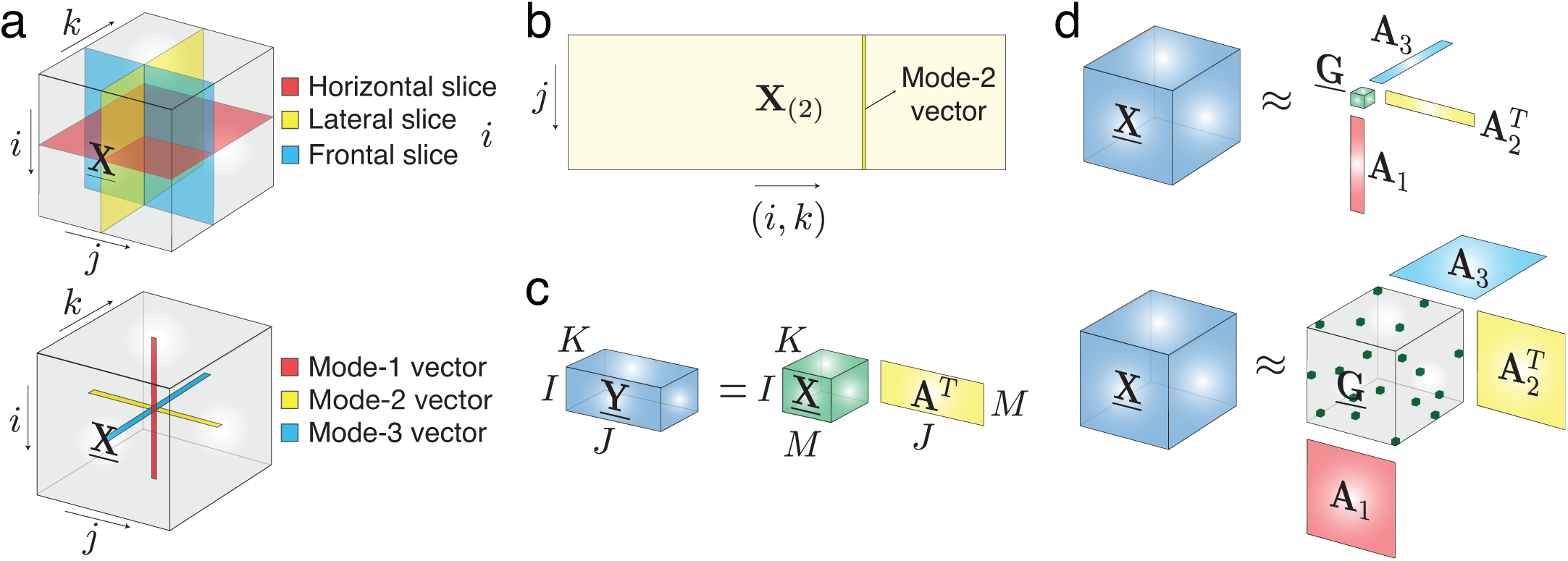
(**a**)Examples of frontal (light blue), lateral (yellow) and horizontal (red) slices of 3D tensor (top),and examples of mode-n vectors (bottom). (**b**) Illustration of the mode-2 unfolded matrix **X**_(2)_. (**c**) Tensor-by-matrix product (example of product in mode-2). (**d**) The classical Tucker decomposition (top;[10]) allows representing a 3D tensor
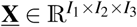 as the product of a core tensor (green) 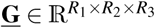 by factor matrices 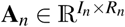 (red, yellow and light blue). Data compression is achieved by considering very small (dense) core tensors <u>**G**</u>, meaning that *R_n_* ≪ *I_n_*. The sparse Tucker Decomposition - SD (bottom; [5]). The core tensor <u>**G**</u> is large but sparse. Data compression is achieved because of the sparsity of the core tensor. See **Supplementary Table 1** for additional information about notation and mathematical definitions.

**Supplementary Fig. 2:**
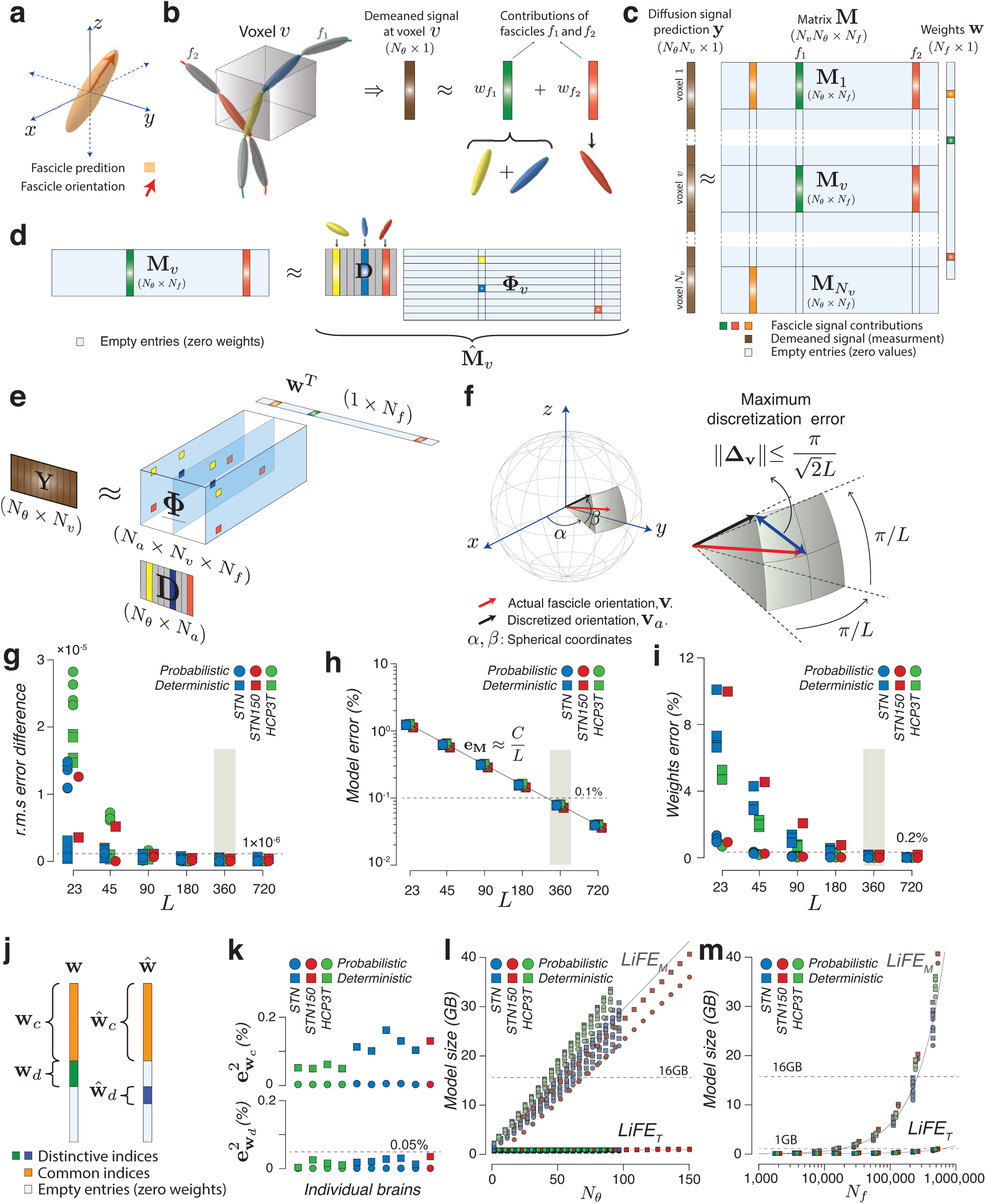
(**a**). A diffusion tensor model whose principal axes are given by the spatial orientation of a fascicle in a voxel **v**. (**b**). Linear model at voxel level: diffusion signal prediction for a voxel intersected by two fascicles, *f*_1_ and *f*_2_, is shown. Fascicle *f*_1_ crosses the voxel from one side to the other and it bends within it, which is modeled as composed by two nodes (yellow and blue). Fascicle *f*_2_ occupies a small portion of the voxel contributing with one diffusion tensor (red). The total prediction signal is the linear combination of the signals associated to existing fascicles within the voxel. (*c*). Linear model for all voxels: LiFE model matrix **M** [9] is obtained by stacking in its columns, the diffusion signal predictions of fascicles. The predicted diffusion signal in all voxels is obtained as **y** ≈ **Mw** where **w** is a sparse vector of weights obtained by solving a convex optimization problem with non-negativity constraints. (**d**). Sparse factorization of the diffusion signal in a voxel: Nonzero columns in block matrix **M_v_** correspond to fascicles passing through *v* (two in our example). Those columns are approximated by combining the diffusion prediction of the dictionary atoms (columns of matrix **D**) with directions closest to the orientation of the fascicles nodes (yellow, blue and red). Non-zero entries in matrix **Φ**_v_ indicate the atoms corresponding to the nodes in the fascicles *f*_1_ and *f*_2_ in the example. (**e**). The LiFE_T_ model: the diffusion signal matrix 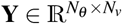 (directions × voxels) is written as a Sparse Tucker Decomposition by using the core Sparse tensor 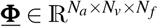, multiplied in mode-1 by the dictionary matrix 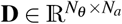 and, in mode-3, by the vector of weights 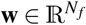. (**f**). Error introduced by the discretization: Left, a regular grid in the sphere is obtained by discretizing the space in spherical coordinates (*α*, *β*). Right, the maximum distance between the fascicle orientation vector *v* and its approximation *v_a_* is inversely proportional to the parameter *L*, i.e. 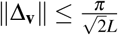. (**g**). Comparison of the global r.m.s error obtained by LiFE and LiFE_T_ models. It is clear that for *L* ≥ 180 LiFE_T_ approximates very well the original LiFE model (difference < 1 × 10 ^−6^). (**h**). Weights error *e_w_* versus parameter *L*. This plot shows that for *L* ≥ 360 LiFE_T_ provides a very good approximation of the LiFE model (*e_w_* < 0.2%). (**i**). The model error *e*_M_ in approximating the matrix **M** with LiFE_T_ is inversely proportional to the parameter *L* as predicted by our theoretical bound (see equation 40). The function 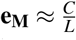 was fitted to the data, which resulted in an estimated value *C* = 28.22, and a fitting error equal to 3.03% (relative root squared error). It is noted that, for *L* ≥ 360, we obtained a model error smaller than 0.1%. (**j**). By defining common (orange) and different (green and blue) subsets of indices within vectors **w** and 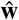 the squared weights error is decomposed as the sum of the squared errors associated to the common and different fascicles (see equation 24). (**k**). The squared error for common fascicles 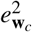 (top) and different fascicles 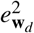 (bottom) computed for STN, STN150 and HCP3T datasets are shown. While the total squared 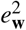 is kept below 0.2%, it is highlighted that it is larger in the deterministic case compared against to the probabilistic case. However, the error associated to the different fascicles is very small compared to the common fascicles, which means that essentially both models use almost the same subset of fascicles. (**l**) and (**m**). Model size (**GB**) scales linearly with the number of directions *N_θ_* and the number of fascicles *N_f_*, however it increases much faster in the LiFE model compared to the LiFE_T_ model.

**Supplementary Fig. 3. (b).**
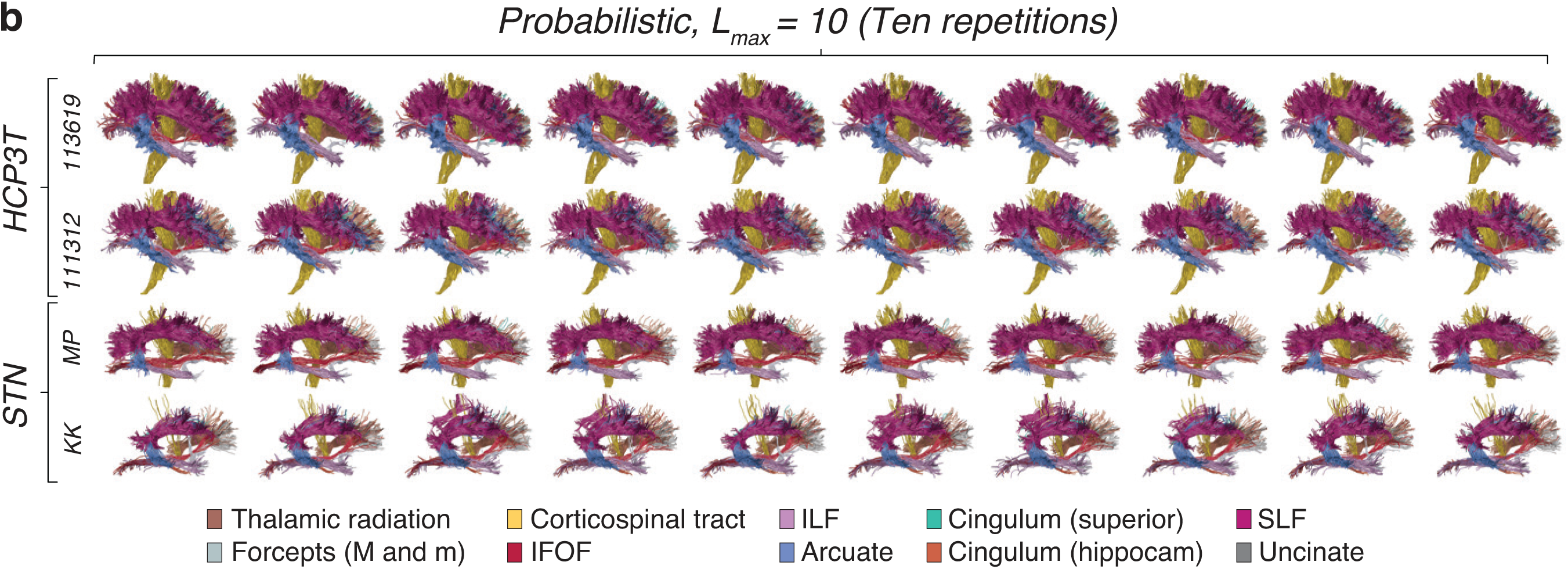
Major human tracts from LiFE optimized connectomes (fascicles with nonzero weights) using Probabilistic tractography (*L_max_* = 10) for two subjects in the STN dataset and two subjects in the HCP3T dataset. Results obtained by repeating the tractography and optimization ten times are shown in different columns. It is highlighted that connectomes are anatomically discriminable across subjects and datasets (rows) but preserving the anatomy among repetitions of the LiFE evaluation (columns).

**Supplementary Fig. 3. (c).**
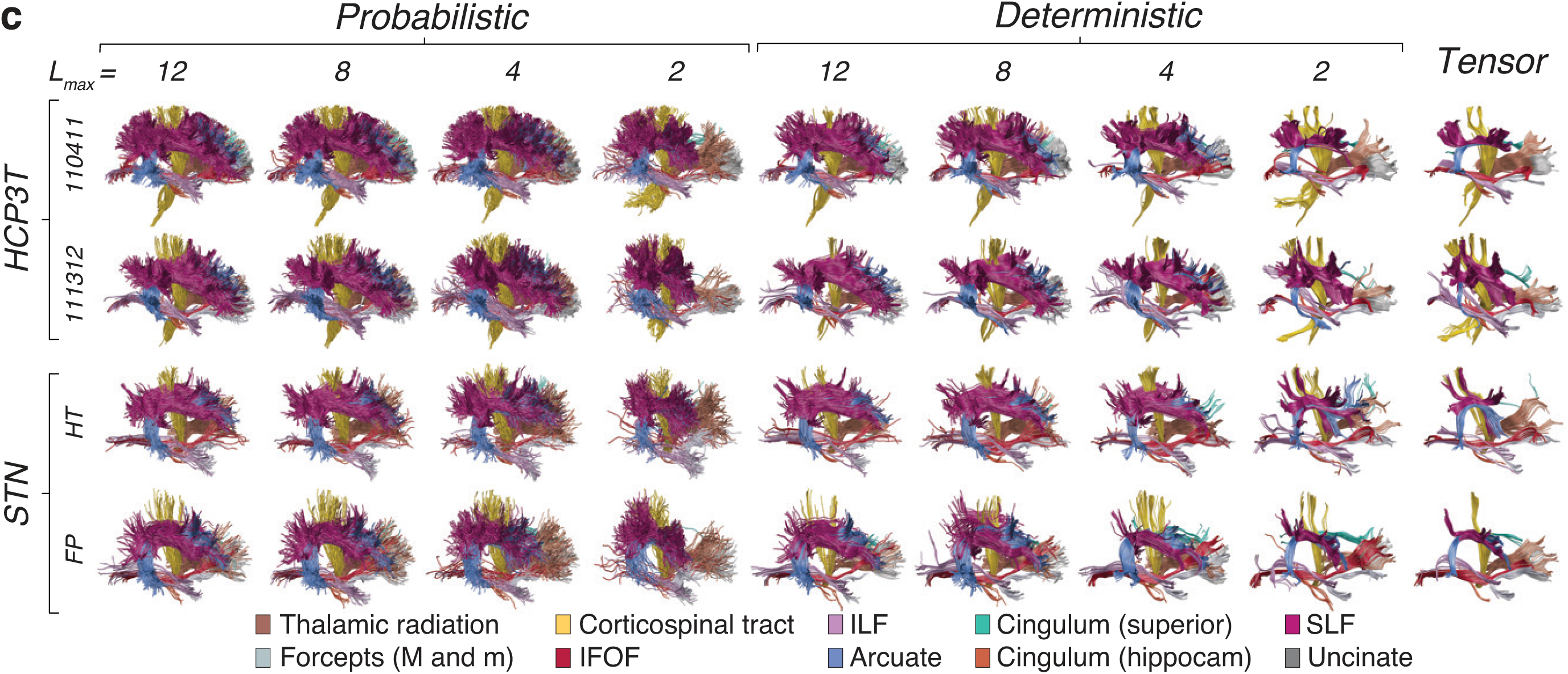
Major human tracts from LiFE optimized connectomes (fascicles with nonzero weights) using probabilistic (*L_max_* = 2,4,6,8,10,12), deterministic(*L_max_* = 2,4,6,8,10,12) and tensor tracking methods for two subjects in the STN dataset and two subjects in the HCP3T dataset. It is highlighted that connectomes are anatomically discriminable across subjects and datasets (rows) and across tracking methods.

**Supplementary Fig. 3. (d).**
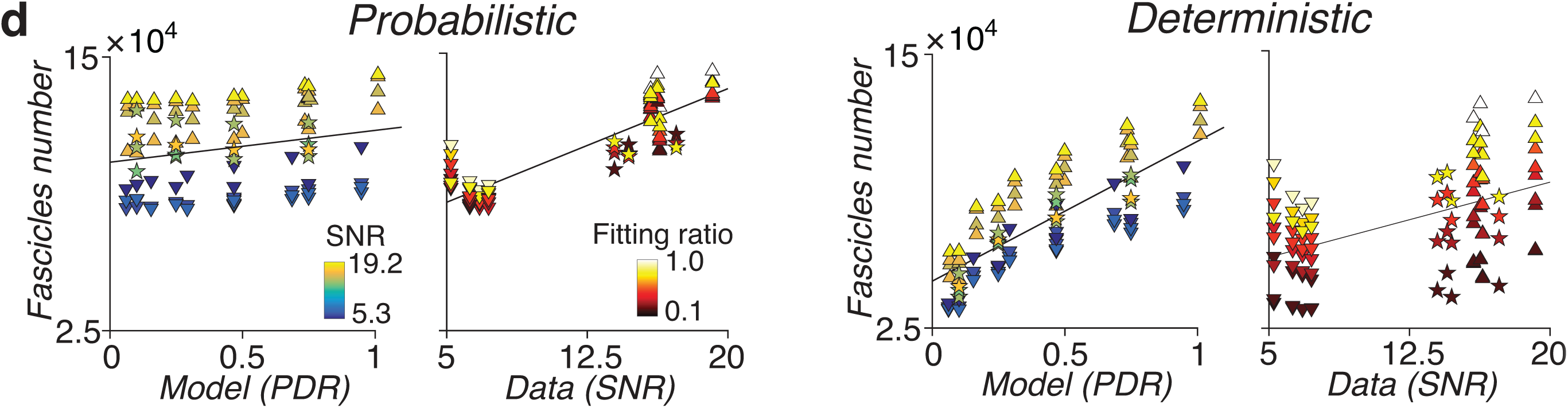
Number of fascicles (nonzero entries in the weights vector) in LiFE optimized connectomes as a function of model quality (*PDR*) and signal quality (*SNR*) for probabilistic (left) and deterministic (right) tracking methods. These plots reveals a clear linear relationship between number of fascicles and *PDR* or *SNR*. A 3D representation of these results is shown in Fig. 3d as plot of number of fascicles versus *PDR* and *SNR*.

**Supplementary Fig. 4.**
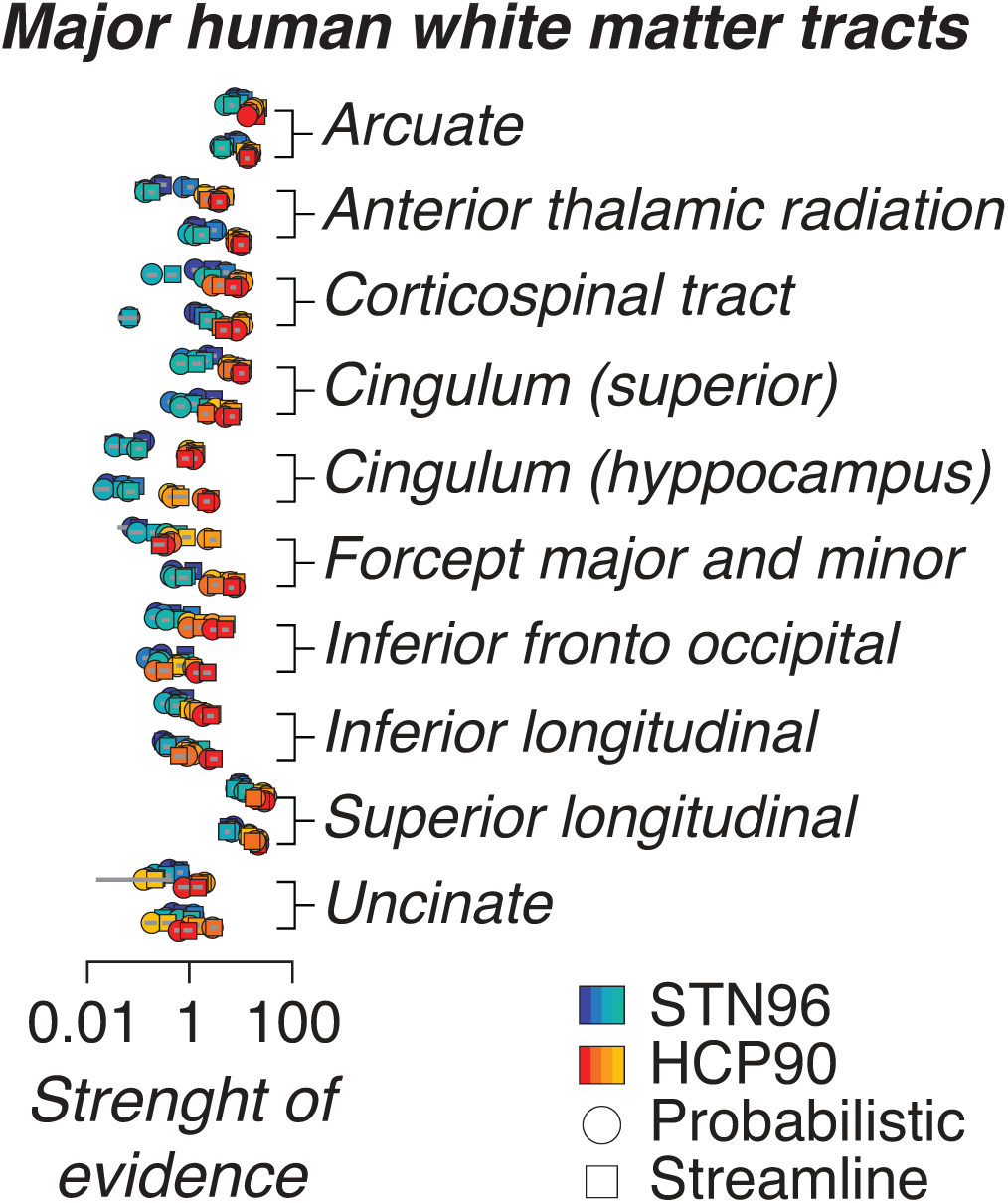
The strength of evidence *S* is a measure of distance between the distributions of r.m.s errors in voxels for the model with virtual lesion and without virtual lesion (see [9] for details). Similarly to the Earth Mover Distance measure (**Fig. 4d**), the strength of evidence is positive for the major tracts, which replicates the statistical evidence of major tracts reported in [9].

**Supplementary Fig. 5.**
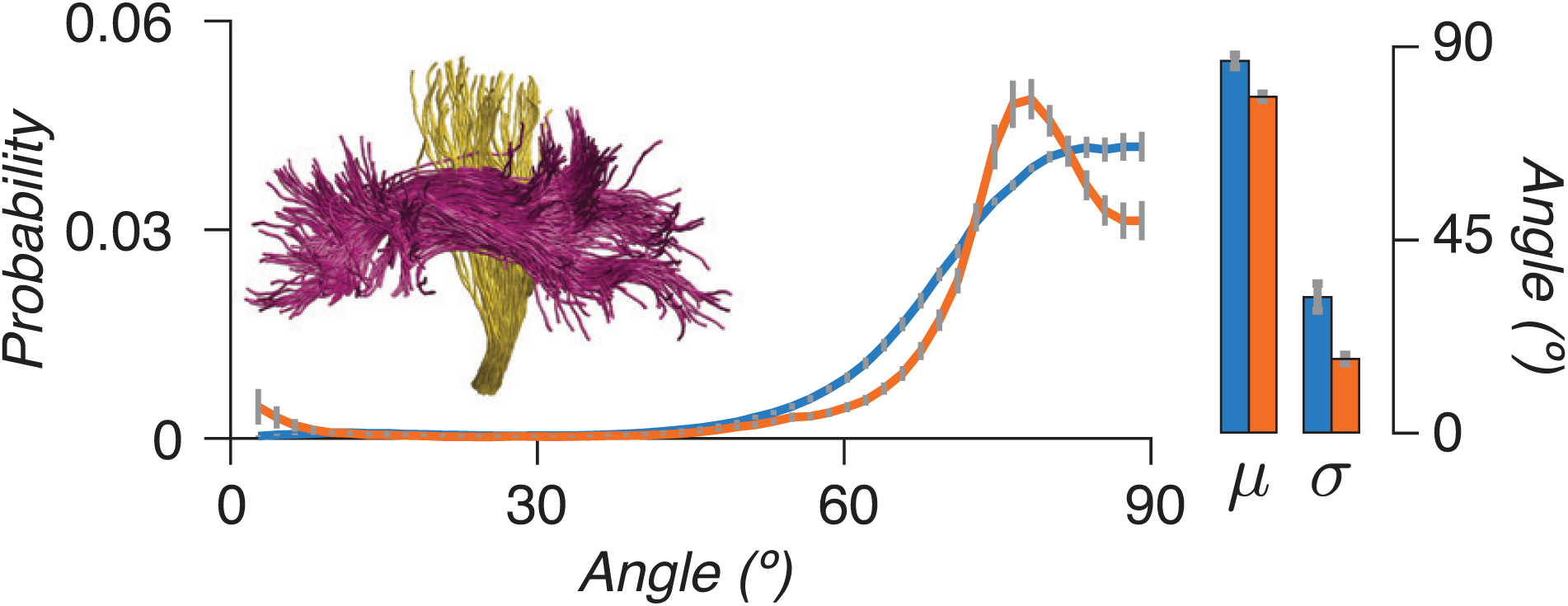
The histograms of angles between fascicles in the Corticospinal tract and SLF tracts are shown for probabilistic (blue) and deterministic (orange) tracking methods with *L_max_* = 10. These distributions show a crossing close to perpendicular similar to the findings for the case of Arcuate and Corticospinal tract in **Fig. 5e**.

**Table 1.** Mathematical notation and definitions for multiway arrays (tensors) and their decomposition.

|  |  |
| --- | --- |
| $\underline{\mathbf{X}}, \mathbf{A}, \mathbf{w}, b$ | A tensor, a matrix, a vector and a scalar |
| $x_{i_1 i_2 \dots i_N}, a_{ij}, w_i$ | Entries of a tensor, a matrix and a vector |
| $\underline{\mathbf{X}}(:, j, k), \underline{\mathbf{X}}(i, :, k), \underline{\mathbf{X}}(i, j, :)$ | Mode-1, mode-2 and mode-3 vectors are obtained by fixing all but one index |
| $\underline{\mathbf{X}}(i, :, :), \underline{\mathbf{X}}(:, j, :), \underline{\mathbf{X}}(:, :, k)$ | Horizontal, lateral and frontal slices are obtained by fixing all but two indices |
| $\mathbf{X}_{(n)} \in \mathbb{R}^{I_n \times I_1 I_2 \dots I_{n-1} I_{n+1} \dots I_N}$ | Mode-$n$ unfolding of tensor $\underline{\mathbf{X}} \in \mathbb{R}^{I_1 \times I_2 \times \dots \times I_N}$ whose entry at row $i_n$ and column $(i_1 - 1)I_2 \dots I_{n-1} I_{n+1} \dots I_N + \dots + (i_{N-1} - 1)I_N + i_N$ is equal to $x_{i_1 i_2 \dots i_N}$ |
| $\underline{\mathbf{Y}} = \underline{\mathbf{X}} \times_n \mathbf{A} \in \mathbb{R}^{I_1 \times \dots \times I_{n-1} \times J \times I_{n+1} \dots \times I_N}$ | tensor by matrix product (in mode-$n$) where $y_{i_1 \dots i_{n-1} j i_{n+1} \dots i_N} = \sum_{i_n=1}^{I_n} x_{i_1 \dots i_n \dots i_N} a_{ji_n}$ |
| $\underline{\mathbf{X}} \approx \underline{\mathbf{G}} \times_1 \mathbf{A}_1 \times_2 \mathbf{A}_2 \times_3 \mathbf{A}_3$ | Tucker decomposition: a 3D tensor $\underline{\mathbf{X}} \in \mathbb{R}^{I_1 \times I_2 \times I_3}$ is represented as the product of a core array $\underline{\mathbf{G}} \in \mathbb{R}^{R_1 \times R_2 \times R_3}$ by factor matrices $\mathbf{A}_n \in \mathbb{R}^{I_n \times R_n}$ |
| $\mathbf{x} = \text{vec}(\underline{\mathbf{X}}) \in \mathbb{R}^{I_1 I_2 \dots I_N}$ | Vectorization of tensor $\underline{\mathbf{X}} \in \mathbb{R}^{I_1 \times I_2 \times \dots \times I_N}$ with the entry at position $i_1 + \sum_{k=2}^N [(i_k - 1)I_1 I_2 \dots I_{k-1}]$ equal to $x_{i_1 i_2 \dots i_N}$ |

## Footnotes

b We note that our experimental results in Fig. 2d showed a compression factor slightly lower than the theoretical estimation because the sparse matrix format implemented in Matlab [7] is relatively more efficient than the sparse multiway format used in the Matlab Tensor Toolbox [2]

